# Microbial signals and lymphotoxin drive TNF-independent death of A20 and ABIN-1 deficient epithelium

**DOI:** 10.1101/2021.11.08.467808

**Authors:** Iulia Rusu, Elvira Mennillo, Zhongmei Li, Jared L. Bain, Xiaofei Sun, Kimberly Ly, Yenny Y. Rosli, Mohammad Naser, Zunqiu Wang, Rommel Advincula, Philip Achacoso, Ling Shao, Bahram Razani, Ophir D. Klein, Alexander Marson, Jessie A. Turnbaugh, Peter J. Turnbaugh, Barbara A. Malynn, Averil Ma, Michael G. Kattah

## Abstract

Anti-TNF antibodies are effective for treating patients with inflammatory bowel disease (IBD), but many patients fail to respond to anti-TNF therapy, highlighting the importance of TNF-independent disease. We previously demonstrated that acute deletion of two IBD susceptibility genes, *A20* (*Tnfaip3*) and *Abin-1* (*Tnip1*), in intestinal epithelial cells (IECs) sensitizes mice to both TNF-dependent and TNF-independent death. Here we show that TNF-independent IEC death after *A20* and *Abin-1* deletion is rescued by germ-free derivation or deletion of *MyD88*, while deletion of *Trif* provides only partial protection. Combined deletion of *Ripk3* and *Casp8*, which inhibits both apoptotic and necroptotic death, completely protects against death after acute deletion of *A20* and *Abin-1* in IECs. *A20* and *Abin-1*-deficient IECs are sensitized to TNF-independent, TNFR-1-mediated death in response to lymphotoxin alpha (LTα) homotrimers. Blockade of LTα *in vivo* reduces weight loss and improves survival when combined with partial deletion of *MyD88*. These data show that microbial signals, *MyD88*, and LTα all contribute to TNF-independent intestinal injury.

**SUMMARY:** Here we show that germ-free derivation, MyD88 deletion, combined Ripk3 and Casp8 deletion, or anti-LTα, all reduce TNF-independent intestinal injury after *A20* and *Abin-1* deletion.

## INTRODUCTION

The intestinal epithelium, a single-cell layer protective lining, is critically important for preventing inflammatory responses to a vast array of microbial stimuli. Inflammatory bowel disease (IBD) is the result of an abnormal immune response to microbial stimuli in genetically susceptible individuals, culminating in intestinal epithelial cell (IEC) injury (1–6). IECs play a central role in IBD pathogenesis, and many candidate IBD-associated genes influence IEC biology, including *ITLN1*, *NOS2, ATG16L1*, *XBP1*, *A20* (*TNFAIP3*), *ABIN-1* (*TNIP1*), among others (7–12). Understanding the genetic, microbial, and environmental factors that influence IEC death and injury may enable identification of biomarkers for precision medicine and highlight novel pathways that could be targeted for treating patients with IBD.

Polymorphisms in *A20* (*TNFAIP3*) and *ABIN-1* (*TNIP1*) are linked to a variety of inflammatory disorders affecting multiple tissues, including IBD (13–17). Germline mutations causing *A20* haploinsufficiency have been identified in patients with a systemic inflammatory disorder characterized in part by intestinal ulcerations, typically with pediatric or even infantile onset (18–21). A20 and ABIN-1 are ubiquitin interacting proteins that interact with each other at the protein level, and both restrict cell death as well as NF-κB signaling downstream of TNF and Toll-like receptors (TLRs) (22–37). A20 and ABIN-1 are both expressed in human and murine intestinal epithelium. Mice with *A20*-deficient IECs develop normally, but are more susceptible to dextran sodium sulfate (DSS)-induced colitis as well as cancer induced by A20-deficient myeloid cells or collaborating oncogenes (38–40). A20 and ABIN-1 have important roles in restricting inflammation in multiple tissue types, but much remains to be learned about the role of A20 and ABIN-1 specifically in intestinal epithelial tissue damage.

We previously demonstrated that (IEC)-specific deletion of either *A20* or *Abin-1* alone does not lead to overt weight loss or intestinal injury, but acute simultaneous deletion of both *A20* and *Abin-1* leads to spontaneous IEC apoptosis, fulminant enterocolitis, and rapid mouse lethality (9). In this setting, *A20* and *Abin-1* cooperatively restrict both TNF-dependent and TNF-independent IEC death. TNF-independent IEC death is substantially less well-characterized than TNF-dependent death, and can involve microbial signals, *Trif* (*Ticam1*), *Zbp1*, interferon signaling, ripoptosome activation, or other inflammatory death pathways (41–46).

Anti-TNF therapy remains one of the most effective approaches for treating Crohn’s disease (CD) and ulcerative colitis (UC), but roughly one-third of patients have no response and one-third of patients lose response over time (47–50). Therefore, understanding the gene products that control TNF-independent IEC injury could have significant translational relevance for anti-TNF non-responders. To better understand the pathways leading to TNF-independent IEC injury we performed *in vivo* and *in vitro* analysis of IECs after acute simultaneous deletion of *A20* and *Abin-1*.

## RESULTS

### Germ-free *A20/Abin-1*^*T-ΔIEC*^*Tnf*^−/−^ mice are protected from TNF-independent IEC death

Mice with floxed A20 (*A20*^*fl/fl*^) and floxed Abin-1 (*Abin-1*^*fl/fl*^) on a *Vil-cre-ER*^*T2*+^ background (*A20/Abin-1*^*T-ΔIEC*^) undergo acute deletion of *A20* and *Abin-1* in IECs upon treatment with tamoxifen, culminating in spontaneous apoptotic IEC death, severe enterocolitis, and rapid mouse lethality (9). This death occurs on a *Tnf*^+/+^ or *Tnf*^−/−^ background, demonstrating the important role of TNF-independent death in this model. Tamoxifen delivery by intraperitoneal (*i.p*.) oil injection has been reported to cause peritoneal inflammation, foam cell formation, and depletion of resident macrophages (51). To exclude the possibility that sterile peritonitis contributes to TNF-independent death in *A20/Abin-1*^*T-ΔIEC*^*Tnf*^−/−^ mice, we treated mice with tamoxifen by oral gavage rather than *i.p*. A higher dose of tamoxifen was required to delete A20 and ABIN-1 in IECs from the small intestine and colon by oral gavage (**Supplemental Figure 1A**), and with this approach *A20/Abin-1*^*T-ΔIEC*^*Tnf*^−/−^ mice died with similar kinetics as *A20/Abin-1*^*T-ΔIEC*^ mice (**Figure 1A**). Enteroids derived from *A20/Abin-1*^*T-ΔIEC*^*Tnf*^−/−^ mice undergo deletion of A20 and ABIN-1 when treated with 200nM 4-hydroxytamoxifen (4-OHT) *in vitro,* but they are protected from spontaneous cell death (**Supplemental Figure 1B-D**). This suggests that IEC-extrinsic factors *in vivo* drive TNF-independent IEC death and mortality in *A20/Abin-1*^*T-ΔIEC*^*Tnf*^−/−^ mice. Since *in vitro* IEC enteroid cultures are sterile, we considered that microbial signals might promote death *in vivo*. While our prior studies suggested that broad spectrum antibiotic treatment was insufficient to rescue *A20/Abin-1*^*T-ΔIEC*^*Tnf*^−/−^ mice (9), we hypothesized that residual microbes in these mice could trigger IEC death in these mice. Accordingly, we derived *A20/Abin-1*^*T-ΔIEC*^*Tnf*^−/−^ germ-free mice by cesarean section. *Tnf*^−/−^ germ-free mice were largely protected from death upon deletion of *A20* and *Abin-1* in IECs (**Figure 1B**). To control for developmental alterations by germ-free derivation, we conventionalized germ-free *A20/Abin-1*^*T-ΔIEC*^*Tnf*^−/−^ mice with cecal contents from corresponding specific-pathogen free (SPF) mice in our facility. Germ-free mice conventionalized with cecal contents from SPF *A20/Abin-1*^*T-ΔIEC*^*Tnf*^*−/−*^ mice (GF-CONV) exhibited rapid mortality upon deletion of *A20* and *Abin-1* (Figure 1B), suggesting the increased survival of germ-free mice was not due to a developmental aberration. Therefore, microbial signals contribute to TNF-independent IEC death in the setting of acute *A20* and *Abin-1* deletion.

**Figure 1.**
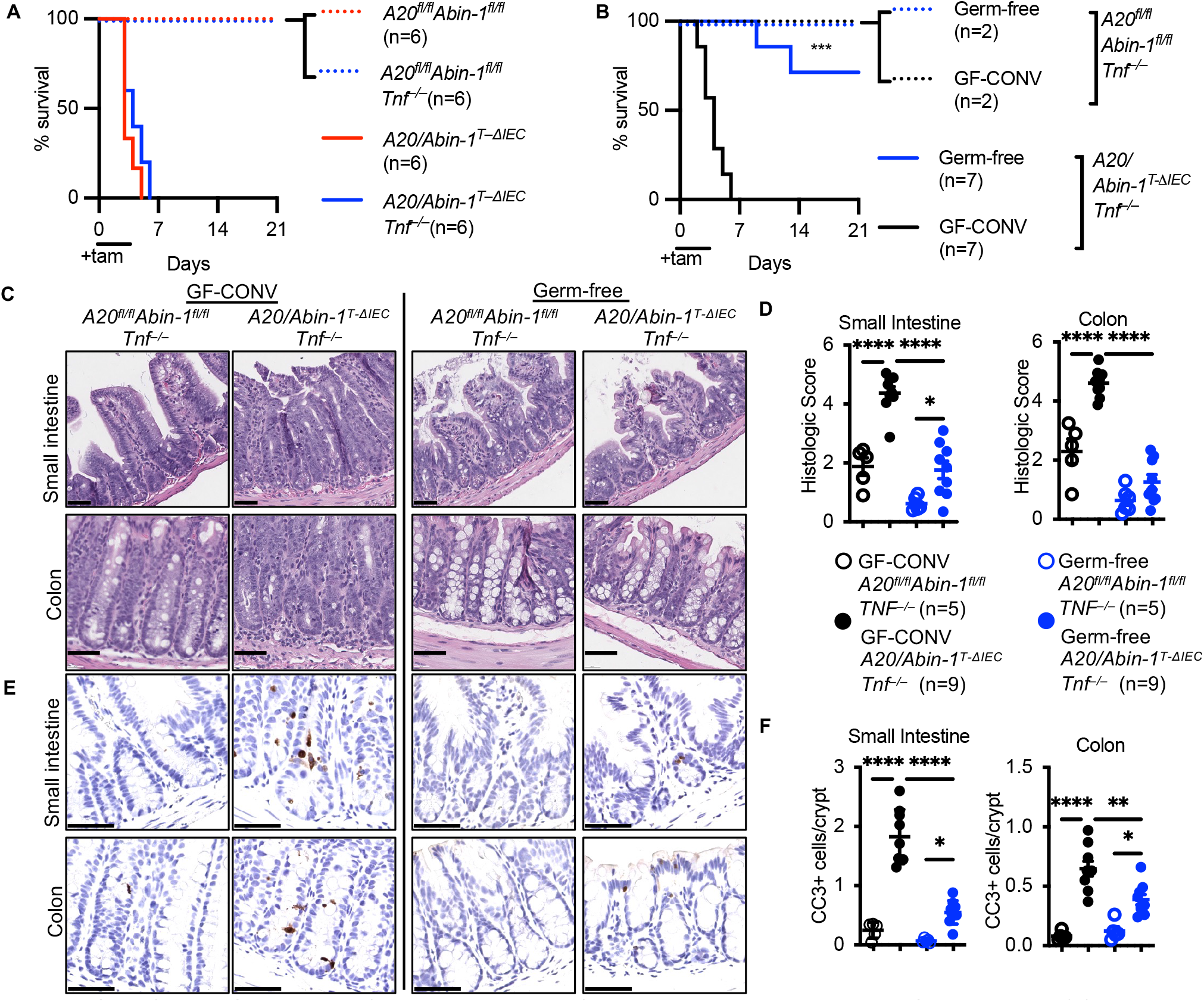
Germ-free *A20/Abin-1*^*T-ΔIEC*^*Tnf*^−/−^ mice are protected from TNF-independent apoptotic IEC death *in vivo*. **(A)** Kaplan-Meier survival curves of the indicated genotypes of tamoxifen-treated mice. (**B**) Kaplan-Meier survival curves of tamoxifen-treated mice with the indicated genotypes, either germ-free or conventionalized with cecal contents from SPF mice (GF-CONV). (**C**) Representative H&E images, (**D**) histological scoring, (**E**) representative CC3 IHC images, and (**F**) CC3+ cells per crypt of small intestine and colon sections 40h after tamoxifen treatment in mice with the indicated genotype; each data point represents one mouse (mean ± SEM). The legend for panel (**F**) is shown in panel (**D**). For panels (**A,B**), statistical significance between *A20/Abin-1*^*T-ΔIEC*^ and *A20/Abin-1*^*T-ΔIEC*^*Tnf*^−/−^ mice was assessed by Log-rank Mantel-Cox test. For panels (**D**) and (**F**), significance was assessed by one-way ANOVA with Tukey’s multiple comparison test. Only significant differences are shown; *=p<0.05, **=p<0.01, ***=p < 0.001; ****=p< 0.0001. Scale bars, 100μm. Data represent at least two independent experiments.

Although germ-free *A20/Abin-1*^*T-ΔIEC*^*Tnf*^−/−^ mice exhibited increased survival, it was unclear whether this was due to reduced IEC death or merely due to broadly reduced septic sequelae under germ-free conditions. Histologically, acute deletion of *A20* and *Abin-1* in the intestinal epithelium caused rapid intestinal epithelial denudation, inflammatory infiltrate, cryptitis, and loss of mucosal architecture in both the small intestine and colon within 40 hours in SPF conventionalized mice (**Figure 1C,D**). In contrast, germ-free *A20/Abin-1*^*T-ΔIEC*^*Tnf*^−/−^ mice exhibited far less histologic injury (**Figure 1C,D**). Since IEC loss in this model is further characterized by massive apoptotic IEC death, we performed cleaved caspase 3 (CC3) immunohistochemistry. In parallel to the reduction in histologic disease severity, we observed dramatically reduced CC3 in IECs of germ-free *A20/Abin-1*^*T-ΔIEC*^*Tnf*^−/−^ mice as compared to SPF conventionalized counterparts (**Figure 1E,F**)

These results highlight that *A20/Abin-1*^*T-ΔIEC*^*Tnf*^−/−^ mice provide a window into studying TNF-independent IEC death. *A20/Abin-1*^*T-ΔIEC*^*Tnf*^−/−^ mice die due to spontaneous fulminant IEC death *in vivo*, but those IECs survive *in vitro* in the absence of hematopoietic cells, autocrine TNF, and microbial stimuli. When microbial stimuli are removed under germ-free conditions, the *A20* and *Abin-1-*deficient IECs survive *in vivo* even with hematopoietic cells and other potential cytotoxic factors present. Interestingly, some germ-free *A20/Abin-1*^*T-ΔIEC*^*Tnf*^−/−^ mice die (Figure 1B), suggesting that there may be some sterile inflammatory factors that can contribute to TNF-independent IEC death, but microbial factors are a primary driver of intestinal inflammation in this model.

### Deletion of *MyD88*, and to a lesser extent *Trif*, rescues *A20/Abin-1*^*T-ΔIEC*^*Tnf*^−/−^ mice

Given the dramatic improvement in survival and intestinal epithelial integrity in germ-free *A20/Abin-1*^*T-ΔIEC*^*Tnf*^−/−^ mice, we hypothesized that microbial signaling through *MyD88* mediated intestinal inflammation in this model. To facilitate combining of multiple mutant alleles, we targeted *MyD88* in *A20/Abin-1*^*T-ΔIEC*^*Tnf*^−/−^ zygotes as previously described (52). Using two guides targeting exon 1 of *MyD88,* we generated two founder strains of mice with deletions at the *MyD88* locus, *A20/Abin-1*^*T-ΔIEC*^*Tnf*^−/−^*MyD88*^−/−^ *C1* and *C2* (**Figure 2A, Supplemental Figure 2A,B**). We confirmed deletion of MYD88 at the protein level in both mouse strains (**Figure 2B**). *A20/Abin-1*^*T-ΔIEC*^*Tnf*^−/−^MyD88^−/−^ *C1* and *C2* behaved identically, and so are presented in aggregate for clarity. Heterozygous deletion of *MyD88* conferred a modest improvement in survival, while complete deletion of *MyD88* led to a marked improvement in survival in *A20/Abin-1*^*T-ΔIEC*^*Tnf*^−/−^ mice (**Figure 2C**). The histologic phenotype paralleled the survival benefit, where the intestinal epithelium from *A20/Abin-1*^*T-ΔIEC*^*Tnf*^−/−^*MyD88*^−/−^ mice exhibited significantly less inflammatory injury in the small intestine and colon as compared *A20/Abin-1*^*T-ΔIEC*^*Tnf*^−/−^ mice (**Figure 2D,E**). Similarly, deletion of *MyD88* significantly reduced the frequency of apoptotic CC3+ IECs as compared to their *MyD88*^+/+^ counterparts (**Figure 2F,G**). *A20/Abin-1*^*T-ΔIEC*^*Tnf*^−/−^*MyD88*^+/−^ mice exhibited intermediate histologic injury and CC3 frequency (**Figure 2D-G**). These results are surprising given that *MyD88* plays a critical role in intestinal homeostasis, and deletion has been reported increase susceptibility to other mouse models of colitis (53–55).

**Figure 2.**
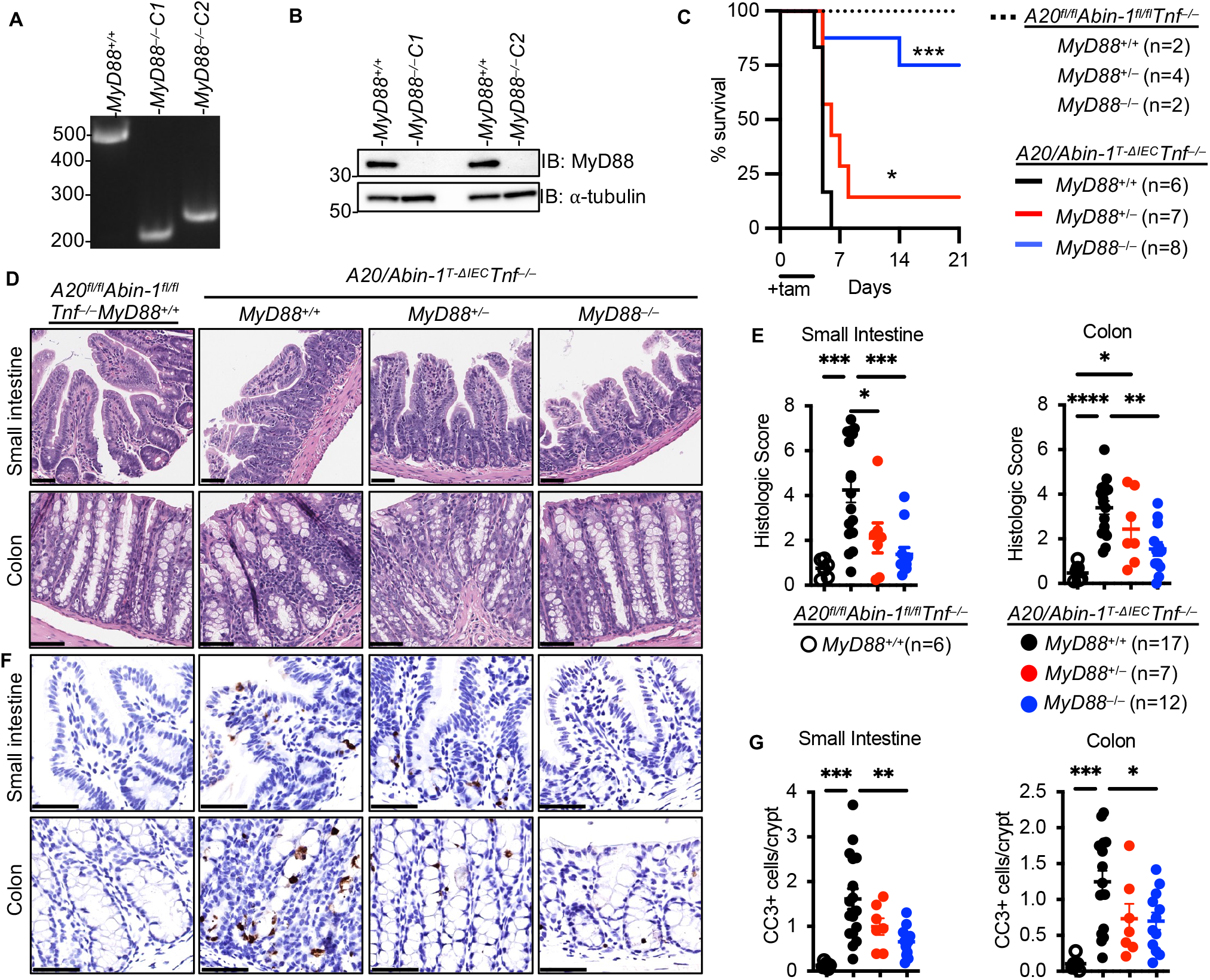
Deletion of *MyD88* significantly improves survival, reduces epithelial injury, and decreases IEC apoptosis in *A20/Abin-1*^*T-ΔIEC*^*Tnf*^−/−^ mice. (**A**) Agarose gel electrophoresis and (**B**) immunoblot of splenocyte lysates from *A20/Abin-1*^*T-ΔIEC*^*Tnf*^−/−^ mice with the indicated *MyD88* genotypes. (**C**). Kaplan-Meier survival curves of the indicated genotypes of tamoxifen-treated mice. (**D**) Representative H&E images, (**E**) histological scoring, (**F**) representative CC3 IHC images, and (**G**) CC3+ cells per crypt of small intestine and colon sections 40h after tamoxifen treatment in mice with the indicated genotype; each data point represents one mouse (mean ± SEM). The legend for panel (**G**) is shown in panel (**E**). For panel (**C**), statistical significance comparing *A20/Abin-1*^*T-ΔIEC*^*Tnf*^−/−^*MyD88*^+/+^ mice to *MyD88*^−/−^ and *MyD88*^+/−^ mice was assessed by Log-rank Mantel-Cox test. For panels (**E**) and (**G**), statistical significance was assessed by one-way ANOVA with Tukey’s multiple comparison test. Only significant differences are shown; *=p<0.05, **=p<0.01, ***=p < 0.001; ****=p< 0.0001. Scale bars, 100μm. Data represent at least two independent experiments.

*MyD88* expression in IECs maintains intestinal epithelial integrity and homeostasis (54, 55). Given the improved survival of *A20/Abin-1*^*T-ΔIEC*^*Tnf*^−/−^*MyD88*^−/−^ mice, we next examined whether MYD88 activation directly induces apoptotic IEC death using *A20/Abin-1*^*T-ΔIEC*^*Tnf*^−/−^ primary small intestinal enteroid cultures. MYD88 mediates signaling downstream of both Toll-like receptors (TLRs) and IL-1 family members (56, 57), so we stimulated IECs with pam3CSK4 (a TLR2/1 agonist). This ligand did not induce significant death in *A20/Abin-1*^*T-ΔIEC*^*Tnf*^−/−^ enteroids (**Figure 3A**). Similarly, IL-1β and IL-18, two IL-1 family members that activate MYD88, did not directly induce death in *A20/Abin-1*^*T-ΔIEC*^*Tnf*^−/−^ enteroids (**Figure 3A, Supplemental Figure 3A**). Although *MyD88* deletion rescues *A20/Abin-1*^*T-ΔIEC*^*Tnf*^−/−^ mice, MYD88 activation was not sufficient for IEC death *in vitro*.

**Figure 3.**
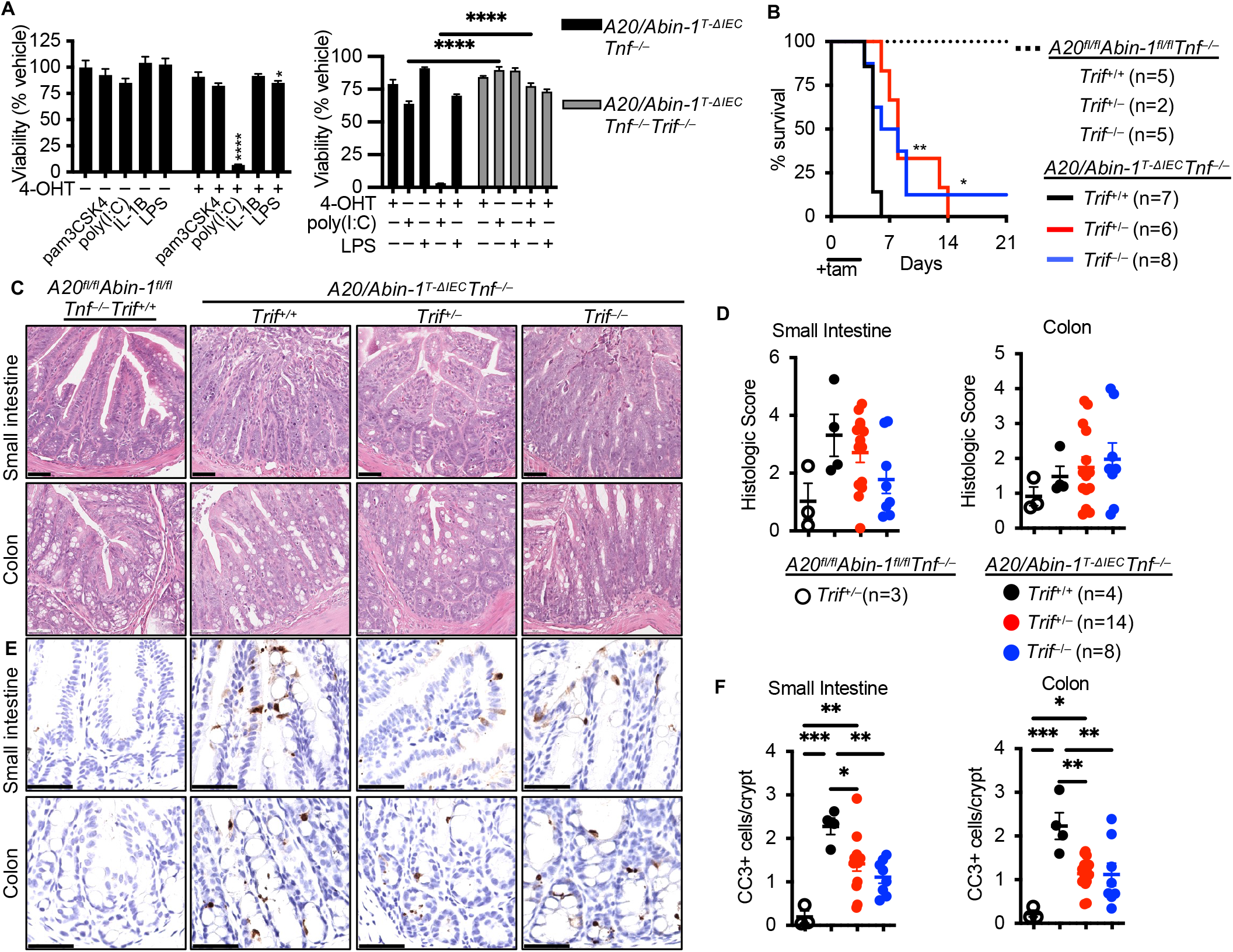
Deletion of *Trif* partially improves survival and modestly reduces IEC apoptosis in *A20/Abin-1*^*T-ΔIEC*^*Tnf*^−/−^ mice. (**A**) Quantitative luminescent cell viability assay of enteroid cultures with the indicated genotype treated with the indicated stimuli (mean ± SEM; pam3CSK4 500ng/ml; poly (I:C) 25ug/ml; IL-1β 25ng/ml; LPS 500ng/ml). (**B**). Kaplan-Meier survival curves of the indicated genotypes of tamoxifen-treated mice. (**C**) Representative H&E images, (**D**) histological scoring, (**E**) representative CC3 IHC images, and (**F**) CC3+ cells per crypt of small intestine and colon sections 40h after tamoxifen treatment in mice with the indicated genotype; each data point represents one mouse (mean ± SEM). The legend for panel (**F**) is shown in panel (**D**). For panel (A), statistical significance was assessed using two-way ANOVA with Bonferroni’s multiple comparison test. For panel (**B**), significance was comparing *A20/Abin-1*^*T-ΔIEC*^*Tnf*^−/−^*Trif*^+/+^ mice to *Trif*^−/−^ and *Trif*^+/−^ mice was assessed by Log-rank Mantel-Cox test. For panels (**D**) and (**F**), statistical significance was assessed by one-way ANOVA with Tukey’s multiple comparison test. Only significant differences are shown; *=p<0.05, **=p<0.01, ***=p < 0.001; ****=p< 0.0001. Scale bars, 100μm. Data represent at least two independent experiments.

TRIF contains a RIP homotypic interaction motif (RHIM) domain and interacts with RIPK3 to induce death signaling downstream of TLR3 in response to the dsRNA analog polyinosinic-polycytidylic acid (poly(I:C)) (44, 58–61). We tested the susceptibility of *A20/Abin-1*^*T-ΔIEC*^*Tnf*^−/−^ enteroids to poly(I:C)-induced death, and we observed that deletion of *A20* and *Abin-1* in IECs dramatically increased susceptibility to poly(I:C)-induced death (**Figure 3A**). LPS, in contrast, induced minimal cytotoxicity (**Figure 3A**). Both poly(I:C) and LPS activate TRIF, but IEC-death in response to LPS has previously been reported to depend on TNF, in contrast to poly(I:C) (44). We deleted *Trif* in *A20/Abin-1*^*T-ΔIEC*^*Tnf*^−/−^ mice using CRISPR/Cas9 editing and generated a strain of mice with a deletion and premature stop codon at the *Trif* locus (**Supplemental Figure 3B,C**). Splenocytes derived from *A20/Abin-1*^*T-ΔIEC*^*Tnf*^−/−^*Trif*^−/−^ mice exhibited significantly reduced IFN-β production in response to poly(I:C), confirming functional deletion at the protein level (**Supplemental Figure 3D**). To determine if *Trif* deletion protected *A20* and *Abin-1* deficient IECs from TNF-independent death *in vitro*, we expanded enteroid cultures from these mice. *A20/Abin-1*^*T-ΔIEC*^*Tnf*^−/^*Trif*^−/−^ enteroids were almost entirely resistant to poly(I:C) and LPS-induced cell death (**Figure 3A**). Therefore, A20 and ABIN-1 cooperatively restrict TRIF-mediated death in response to TLR3 and TLR4 agonists *in vitro*.

Given that *Trif* deletion rescued TNF-independent death in response to poly(I:C) and LPS *in vitro*, we treated *A20/Abin-1*^*T-ΔIEC*^*Tnf*^−/−^*Trif*^−/−^ mice with tamoxifen to see if *Trif* deletion also protected *A20* and *Abin-1* deficient IECs *in vivo*. Deletion of *Trif* in *A20/Abin-1*^*T-ΔIEC*^*Tnf*^−/−^ mice provided a modest survival benefit, with slightly delayed median survival relative to *Trif*^+/+^ controls (**Figure 3B**). Interestingly, heterozygous and homozygous deletion of *Trif* conferred similar survival benefit. We examined the small intestine and colon at 40h after tamoxifen-induced deletion of *A20* and *Abin-1* in *A20/Abin-1*^*T-ΔIEC*^*Tnf*^−/−^*Trif*^−/−^ mice and saw a trend toward improvement in histologic severity, but this was not statistically significant when compared to the *A20/Abin-1*^*T-ΔIEC*^*Tnf*^−/−^ mice (**Figure 3C,D**). Deletion of *Trif* did partially reduce CC3 frequency to some extent, but not enough to prevent lethality in the *A20/Abin-1*^*T-ΔIEC*^*Tnf*^−/−^ mice (**Figure 3E,F**). While poly(I:C) potently induces *Trif*-dependent cytotoxicity in *A20* and *Abin-1*-deficient IECs *in vitro,* deletion of *Trif* does not protect against TNF-independent intestinal injury *in vivo*.

### Combined deletion of *Ripk3* and *Casp8* completely protects *A20/Abin-1*^*T-ΔIEC*^ IECs from death

Deletion of *Trif* did not reverse apoptotic IEC death in *A20/Abin-1*^*T-ΔIEC*^*Tnf*^−/−^ mice, but A20 is known to inhibit necroptosis as well as apoptosis (24, 62), and combined deletion of *A20* and *Abin-1* sensitizes IECs to both Casp8-dependent apoptotic death and RIPK3-dependent necroptotic death *in vitro* (9). Deletion of *Ripk3* does not prevent IEC injury after acute deletion of *A20* and *Abin-1*, so we hypothesized that combined deletion of *Ripk3* and *Casp8* would rescue *A20/Abin-1*^*T-ΔIEC*^ mice. We performed CRISPR/Cas9 editing of the *Casp8* locus in *A20/Abin-1*^*T-ΔIEC*^*Ripk3*^−/−^ mice. This yielded a strain of mice with a premature stop codon in exon 2 of *Casp8* (**Supplemental Figure 4A**), with deletion of CASP8 at the protein level confirmed by immunoblot (**Supplemental Figure 4B**). *A20/Abin-1*^*T-ΔIEC*^*Ripk3*^−/−^ mice exhibited 100% lethality, but *A20/Abin-1*^*T-ΔIEC*^*Ripk3*^−/−^ *Casp8*^−/−^ mice were completely protected from death after treatment with tamoxifen (**Figure 4A**). Histologically, *A20/Abin-1*^*T-ΔIEC*^*Ripk3*^−/−^*Casp8*^−/−^ mice exhibited markedly improved histologic disease severity (**Figure 4B,C**) and CC3 frequency was reduced to near background levels (**Figure 4D,E**). *In vitro*, enteroids derived from *A20/Abin-1*^*T-ΔIEC*^*Ripk3*^−/−^*Casp8*^−/−^ mice were completely protected from both spontaneous IEC death (**Figure 4F**) and TRIF-mediated cell death (**Supplemental Figure 4C**). These data collectively demonstrate that simultaneous suppression of both Casp8-dependent apoptosis and RIPK3-dependent necroptosis completely preserves IEC survival after acute deletion of *A20* and *Abin-1 in vitro* and *in vivo*.

**Figure 4.**
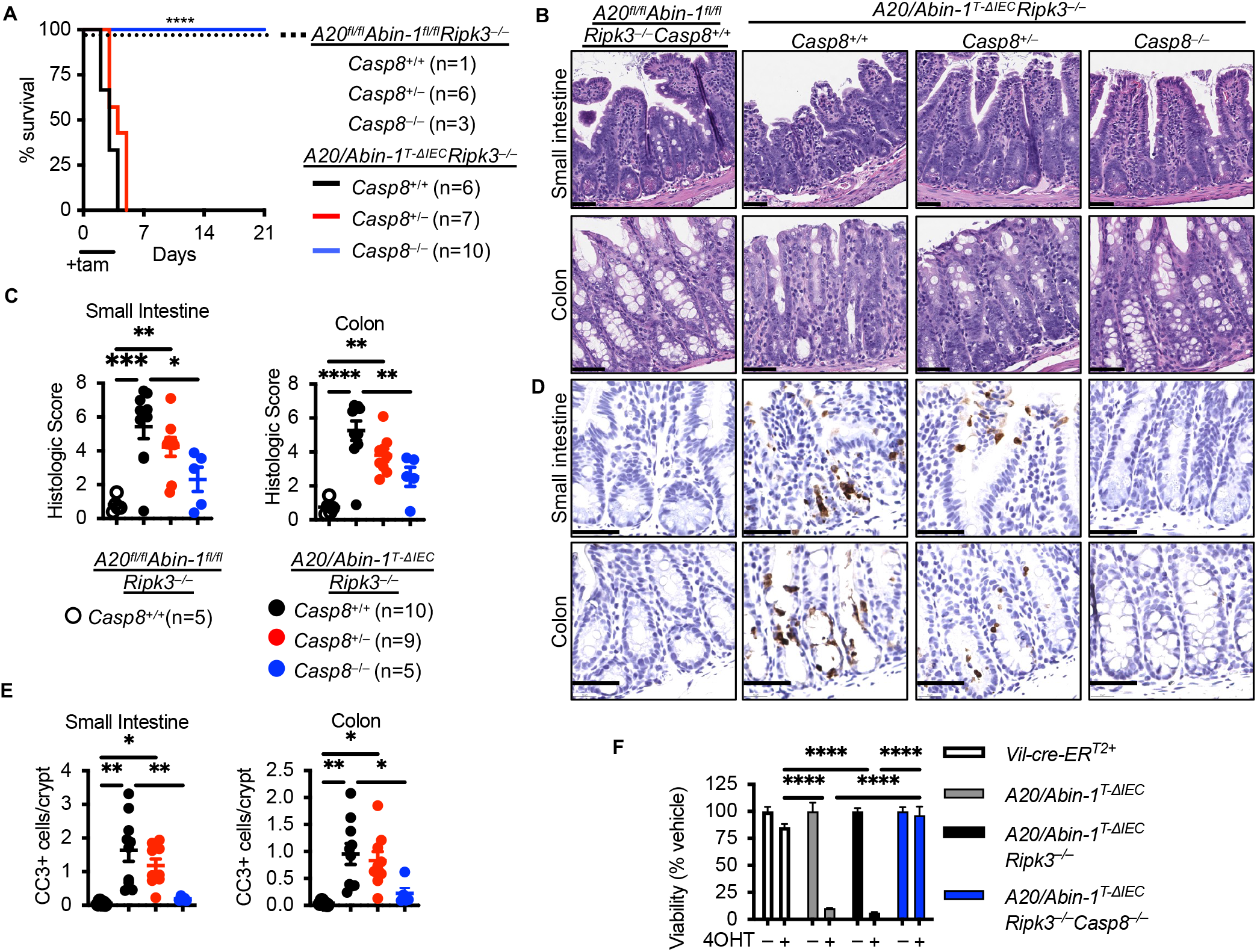
Combined deletion of *Ripk3 and Casp8* completely protects against death, epithelial injury, and IEC apoptosis in *A20/Abin-1*^*T-ΔIEC*^ mice. (**A**). Kaplan-Meier survival curves of the indicated genotypes of tamoxifen-treated mice. (**B**) Representative H&E images, (**C**) histological scoring, (**D**) representative CC3 IHC images, and (**F**) CC3+ cells per crypt of small intestine and colon sections 40h after tamoxifen treatment in mice with the indicated genotype; each data point represents one mouse (mean ± SEM). The legend for panel (**E**) is shown in panel (**C**). (**F**) Quantitative luminescent cell viability assay of enteroids with the indicated genotype treated with indicated stimuli (mean ± SEM). For panel (**A**), statistical significance comparing *A20/Abin-1*^*T-ΔIEC*^*Ripk3*^−/−^*Casp8*^+/+^ mice to *Casp8*^+/–^ and *Casp8*^−/−^ mice was assessed by Log-rank Mantel-Cox test. For panels (**C**), (**E**), and (**F**), statistical significance was assessed by one-way ANOVA with Tukey’s multiple comparison test. Only significant differences are shown; *=p<0.05, **=p<0.01, ***=p < 0.001; ****=p< 0.0001. Scale bars, 100μm. Data represent at least two independent experiments.

### LTα_3_ induces apoptosis and necroptosis downstream of TNFR1 in *A20/Abin-1*^*T-ΔIEC*^*Tnf*^−/−^ enteroids

As combined deletion of *Ripk3* and *Casp8* completely prevented IEC death after deletion of *A20* and *Abin-1*, but *Trif* deletion did not provide equivalent survival benefit, we hypothesized that another TNF superfamily member may be contributing to TNF-independent, IEC-extrinsic death *in vivo.* We previously reported that the TNF superfamily members TNF-like weak inducer of apoptosis (TWEAK), Fas ligand, and TNF-related apoptosis inducing ligand (TRAIL) failed to induce significant death of *A20/Abin-1*^*T-ΔIEC*^*Tnf*^−/−^ enteroids *in vitro* (9). Here we tested receptor activator of nuclear factor kappa-Β ligand (RANKL), “homologous to lymphotoxin, exhibits inducible expression and competes with HSV glycoprotein D for binding to herpesvirus entry mediator, a receptor expressed on T lymphocytes” (LIGHT), and TNF-like cytokine 1A (TL1A), none of which induced significant IEC death (**Figure 5A**).

**Figure 5.**
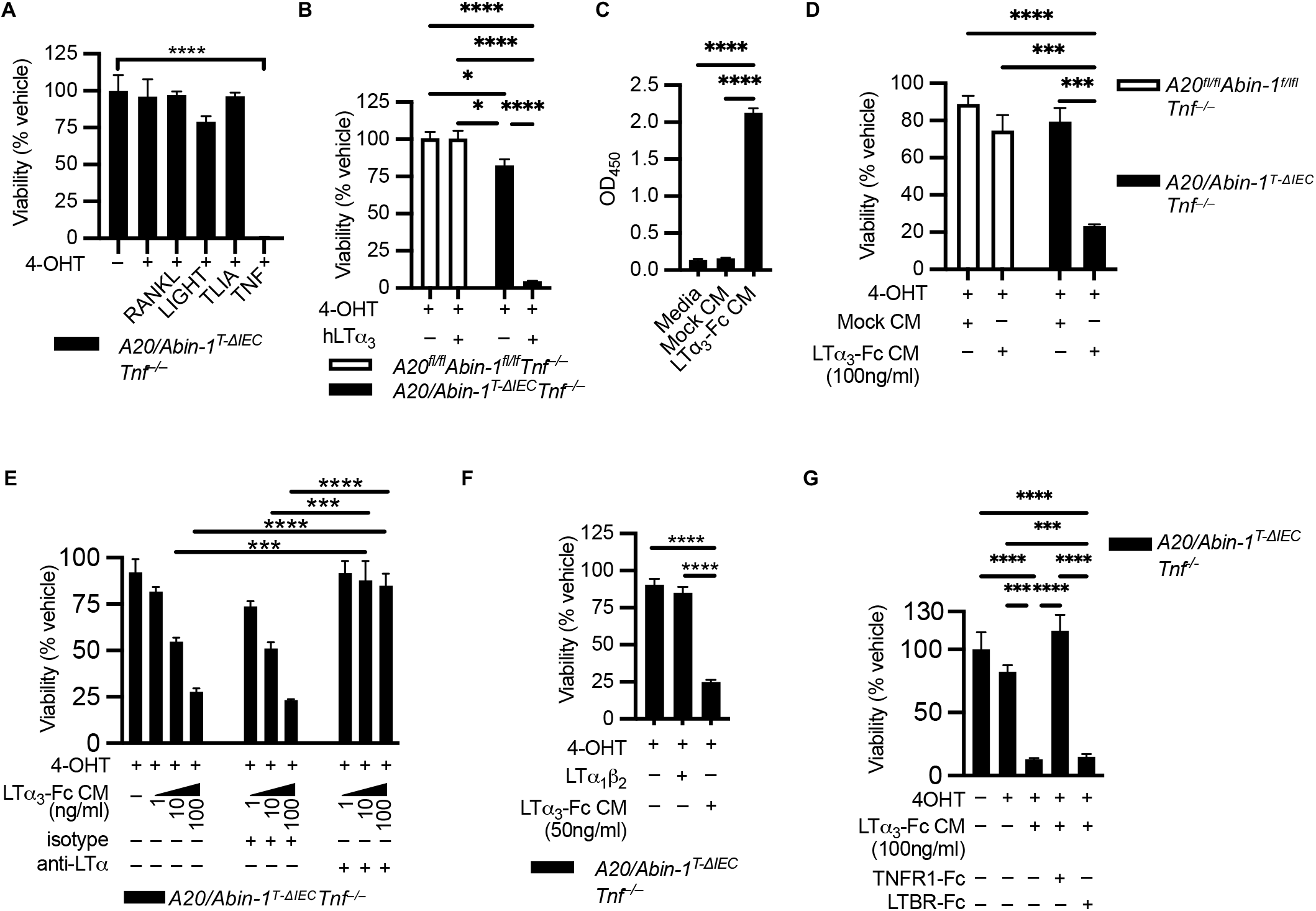
LTα_3_ induces death in *A20/Abin-1*^*T-ΔIEC*^*Tnf*^−/−^ enteroids through activation of TNFR1. (**A,B,D-G**) Quantitative luminescent cell viability assay of enteroids with the indicated genotype treated with indicated stimuli (mean ± SEM; RANKL, LIGHT, TL1A, TNF 25ng/ml; hLTα_3_ 5ng/ml; LTα_1_β_2_ 50ng/ml; anti-LTα and isotype control 10ug/ml; TNFR1-Fc, LTBR-Fc 100ug/ml). (**C**) ELISA of undiluted conditioned media (CM) from 293T cells either mock transduced (Mock CM) or stably transduced with a mouse LTα_3_-Fc expression construct (LTα_3_-Fc CM) with anti-mIgG2a capture and anti-LTα detection antibodies (mean ± SEM). For panel (**A**), significance was assessed by one-way ANOVA with Dunnett’s multiple comparison test relative to vehicle. For panel (**B, E**), significance was assessed using two-way ANOVA with Tukey’s multiple comparison test. For panels (**C,D,F,G**), significance was assessed using one-way ANOVA with Tukey’s multiple comparison test. Only significant differences are shown; *=p<0.05, **=p<0.01, ***=p < 0.001; ****=p< 0.0001. Data represent at least two independent experiments.

Although none of those alternative TNF superfamily ligands induced death in *A20/Abin-1*^*T-ΔIEC*^*Tnf*^−/−^ enteroids, we previously showed that deletion of *A20* and *Abin-1* increases sensitivity of the TNF receptor 1 (TNFR1) to TNF by 1000-fold in IECs (9). Given that TNFR1 sensitivity is significantly increased in *A20* and *Abin-1-* deficient IECs, we considered alternative TNFR1 ligands. Lymphotoxin alpha homotrimers (LTα_3_) are reported to bind to TNFR1 and activate death signaling (63–65). We tested human LTα_3_ and observed significant increased susceptibility to death in *A20* and *Abin-1-*deficient IECs (**Figure 5B**). To test mouse LTα_3_, we generated a cell line constitutively expressing mouse LTα_3_-Fc fusion protein and validated the presence of soluble fusion protein (**Figure 5C**). LTα_3_-Fc fusion protein conditioned media (LTα3-Fc CM) induced significant death in *A20/Abin-1*^*T-ΔIEC*^*Tnf*^−/−^ enteroids (**Figure 5D**). Since LTα_3_-Fc CM could contain other factors that drive IEC death *in vitro*, we wanted to determine if a LTα monoclonal antibody reversed this activity *in vitro*. A LTα-specific monoclonal antibody was previously developed and reported to inhibit collagen-induced arthritis (CIA) (66). We generated purified monoclonal antibody from this hybridoma and observed complete prevention of *A20/Abin-1*^*T-ΔIEC*^*Tnf*^−/−^ IEC death in response LTα_3_-Fc CM *in vitro* (**Figure 5E**). As a negative control, an isotype control antibody did not block cytotoxic activity (**Figure 5E**). Subsequently, we wanted to determine if TNF-independent IEC death in *A20* and *Abin-1*-deficient IECs in response to LTα_3_-Fc CM was indeed mediated by TNFR1. We added recombinant mouse LTα_1_β_2_, which binds to LTBR but not TNFR1 (64), and we did not observe any cytotoxicity (**Figure 5F**). This argues that deletion of *A20* and *Abin-1* in IECs does not increase susceptibility to LTBR-mediated death. We added recombinant mouse LTBR-Fc and TNFR1-Fc fusion proteins and observed that only TNFR1-Fc completely protected *A20/Abin-1*^*T-ΔIEC*^*Tnf*^−/−^ enteroids from death (**Figure 5G**). These data demonstrate that deletion of *A20* and *Abin-1* in IECs unveils a profound sensitivity to LTα_3_-induced death downstream of TNFR1. It is important to highlight that the death assays we performed used primary, non-immortalized IECs in the absence of any death-sensitizing agents (e.g., cycloheximide), unlike most prior studies investigating TNF and LTα_3_-induced death (Chaturvedi et al., 1994; Etemadi et al., 2013).

Having determined that A20 and ABIN-1-deficient IECs are sensitized to LTα_3_-induced death downstream of TNFR1, we wanted to ascertain if LTα_3_-induced death was primarily apoptotic or necroptotic. Receptor interacting serine/threonine kinase 1 (RIPK1) binds caspase 8 and RIPK3, and RIPK1’s kinase activity can support both apoptotic and necroptotic death (43, 45, 67, 68). Necrostatin-1s (Nec1s) is a small molecule RIPK1 kinase inhibitor that suppresses necroptosis and partially suppresses apoptosis (9, 69). As expected, the addition of Nec1s partially prevented LTα_3_-induced death in *A20/Abin-1*^*T-ΔIEC*^*Tnf*^−/−^ enteroids (**Figure 6A**). Caspase inhibition increases RIPK3-dependent necroptosis (70–72). Addition of emricasan, a pharmacologic pan-caspase inhibitor, caused increased death in response to LTα_3_, indicating that *A20/Abin-1*^*T-ΔIEC*^*Tnf*^−/−^ enteroids are sensitized to LTα_3_-induced necroptosis. In agreement with these *in vitro* death assays, deletion of A20 and ABIN-1 in *A20/Abin-1*^*T-ΔIEC*^*Tnf*^−/−^ enteroids leads to significantly increased CC3, cleaved caspase 8 (CC8), and cleaved PARP in response to LTα_3_ as compared to control *A20*^*fl/fl*^*Abin-1*^*fl/fl*^*Tnf*^−/−^ enteroids (**Figure 6B**). There was no significant increase in phosphorylated RIPK3 (pRIPK3) (**Figure 6B**). This pattern is consistent with apoptosis as the primary mode of death in *A20/Abin-1*^*T-ΔIEC*^*Tnf*^−/−^ enteroids in response to LTα_3_. The addition of Nec1s partially reduced CC3 and CC8 (**Figure 6B**), which is consistent with partial inhibition of apoptosis in response to RIPK1 kinase activity inhibition. The addition of emricasan reduced CC3 and CC8 but markedly increased pRIPK1 and pRIPK3, consistent with increased LTα_3_-induced necroptotic death when caspase activity is inhibited (**Figure 6B**). These data demonstrate that combined deletion of *A20* and *Abin-1* sensitizes IECs to both TNF-independent apoptosis and necroptosis in response to LTα_3_, although apoptosis is the dominant death pathway.

**Figure 6.**
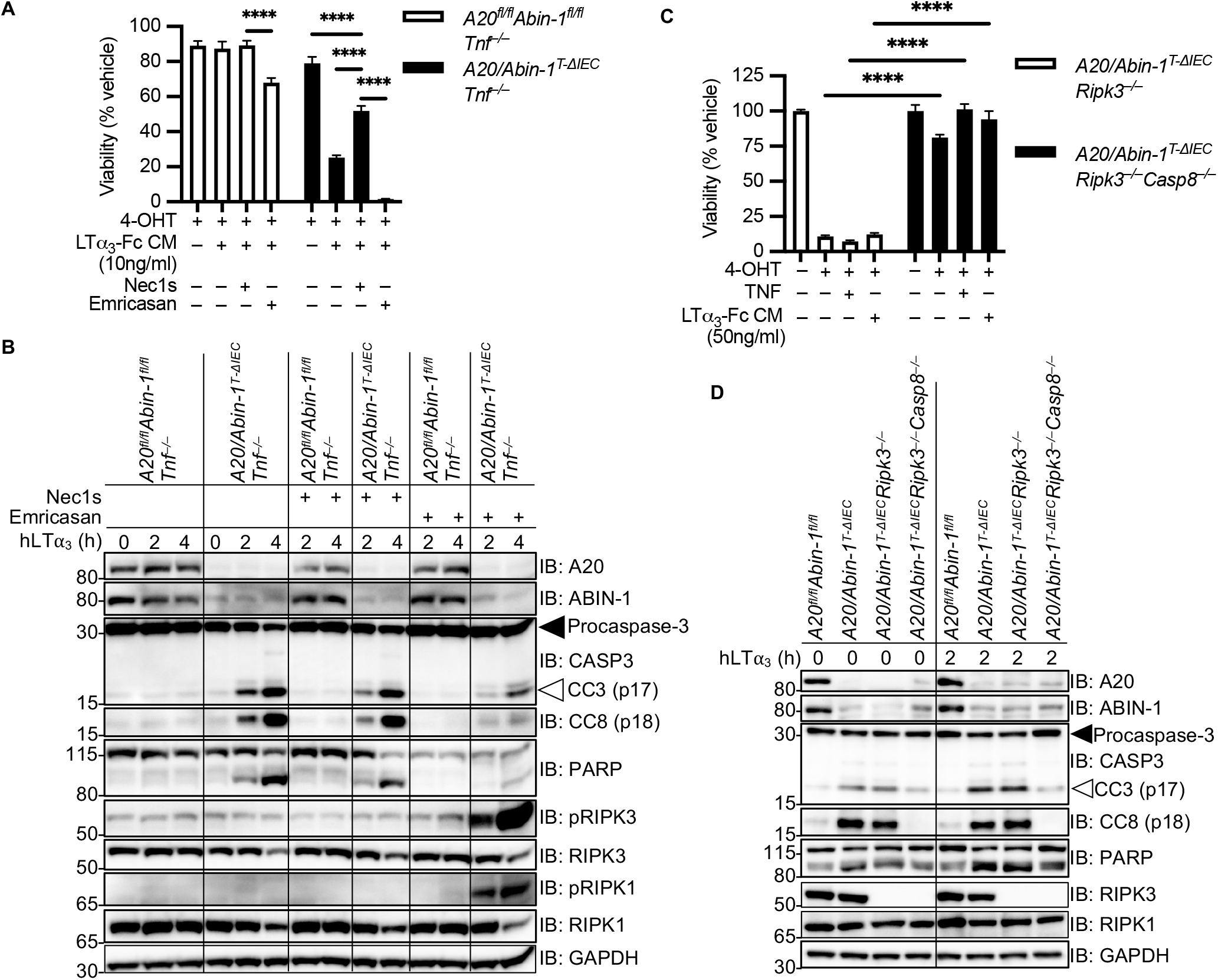
LTα_3_ can induce both Casp8-dependent apoptosis and Ripk3-dependent necroptosis in *A20/Abin-1*^*T-ΔIEC*^*Tnf*^−/−^ enteroids. (**A,C**) Quantitative luminescent cell viability assay of enteroids with the indicated genotypes treated with indicated stimuli (mean ± SEM; Nec1s, Emricasan 50μM; TNF 50ng/ml). (**B,D**) Immunoblotting analyses of enteroid cultures with the indicated genotypes treated with 4-OHT for 24h followed by recombinant hLTα_3_ 20ng/ml for the indicated time, with or without 50μM Nec1s or 50μM emricasan as indicated. Lysates were immunoblotted with the antibodies indicated on the right. Solid arrows indicate full-length protein; open arrows indicate cleaved protein. For panel (**A**), significance was assessed using two-way ANOVA with Dunnett’s multiple comparison test relative to LTα_3_-Fc CM. For panel (**C**), significance was assessed by two-way ANOVA with Bonferroni’s multiple comparison test comparing between genotypes for each stimulation condition. Only significant differences are shown; *=p<0.05, **=p<0.01, ***=p < 0.001; ****=p< 0.0001. Data represent at least two independent experiments.

To further examine necroptosis and apoptosis, we stimulated enteroids derived from *A20/Abin-1*^*T-ΔIEC*^*Ripk3*^−/−^ and *A20/Abin-1*^*T-ΔIEC*^*Ripk3*^−/−^*Casp8*^−/−^ mice with TNF and LTα_3_. The addition of exogenous TNF or LTα_3_ did not induce any additional cytotoxicity in *A20/Abin-1*^*T-ΔIEC*^*Ripk3*^−/−^*Casp8*^−/−^ enteroids (**Figure 6C**). These *in vitro* death assays demonstrate that simultaneous blockade of RIPK3-dependent necroptosis and CASP8-dependent apoptosis is required to rescue A20 and ABIN-1-deficient IECs from death in response to TNF or LTα_3_. We next examined death signaling in *A20/Abin-1*^*T-ΔIEC*^*Ripk3*^−/−^ and *A20/Abin-1*^*T-ΔIEC*^*Ripk3*^−/−^*Casp8*^−/−^ enteroids. Consistent with our prior studies, *A20/Abin-1*^*T-ΔIEC*^*Ripk3*^−/−^ enteroids demonstrated increased spontaneous CC3 and CC8 upon deletion of *A20* and *Abin-1*, as compared to *A20*^*fl/fl*^*Abin-1*^*fl/fl*^ enteroids (Figure 6D). *A20/Abin-1*^*T-ΔIEC*^*Ripk3*^−/−^*Casp8*^−/−^ enteroids, in contrast, did not exhibit CC3, CC8, or cleaved PARP relative to *A20*^*fl/fl*^*Abin-1*^*fl/fl*^ enteroids, even in the presence of exogenous LTα_3_ (**Figure 6D**). Therefore, combined deletion of RIPK3-dependent necroptosis and CASP8-dependent apoptosis completely protects *A20/Abin-1*^*T-ΔIEC*^ IECs from TNF- or LTα_3_-induced apoptotic and necroptotic death downstream of TNFR1.

### LTα blockade combined with partial deletion of *MyD88* protects against TNF-independent death in *A20/Abin-1*^*T-ΔIEC*^*Tnf*^−/−^ mice

To determine if LTα_3_ contributes to IEC injury *in vivo*, we first performed chromogenic RNA-*in situ* hybridization (RNA-ISH) for *Lta* in the intestine after IEC-deletion of *A20* and *Abin-1*. *A20/Abin-1*^*T-ΔIEC*^*Tnf*^−/−^ mice exhibited increased *Lta* positive cells in both the small intestine and colon as compared to *A20*^*fl/fl*^*Abin-1*^*fl/fl*^*Tnf*^−/−^ mice, correlating increased *Lta* frequency with histologic severity upon deletion of *A20* and *Abin-1* in IECs (**Figure 7A,B**). Similarly, qPCR analysis of the small intestine and colon demonstrated increased *Lta* mRNA in *A20/Abin-1*^*T-ΔIEC*^*Tnf*^−/−^ mice as compared to *A20*^*fl/fl*^*Abin-1*^*fl/fl*^*Tnf*^−/−^, *A20/Abin-1*^*T-ΔIEC*^*Tnf*^−/−^*MyD88*^−/−^, and germ-free *A20/Abin-1*^*T-ΔIEC*^*Tnf*^−/−^ mice (**Figure 7C**). The increased *Lta* mRNA by RNA-ISH and qPCR paralleled the pattern of intestinal injury we observed in mice with these genotypes.

**Figure 7.**
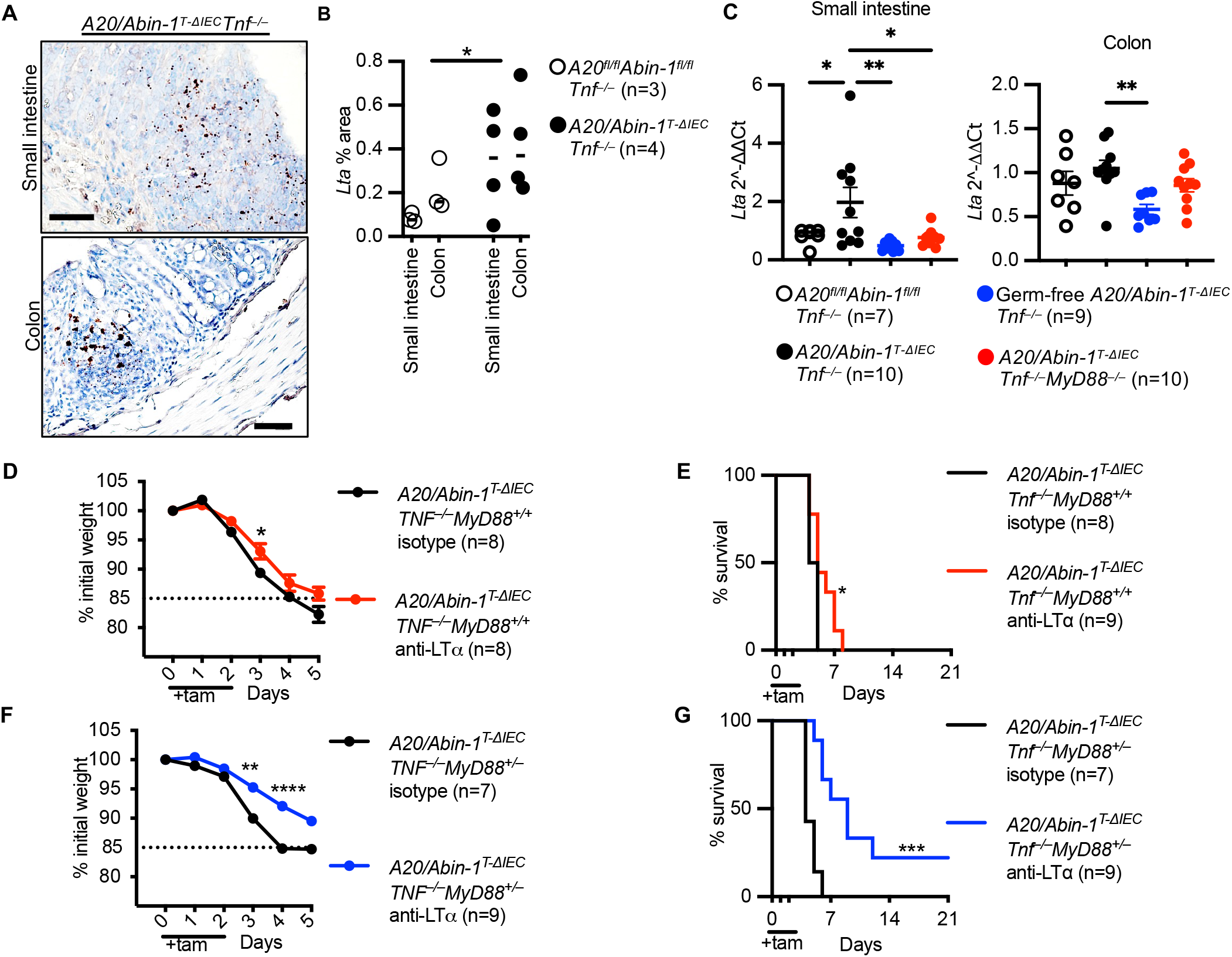
Intestinal *Lta* is increased in *A20/Abin-1*^*T-ΔIEC*^*Tnf*^−/−^ mice and anti-LTα improves survival in *A20/Abin-1*^*T-ΔIEC*^*Tnf*^−/−^*MyD88*^+/+^ and *MyD88*^+/–^ mice. (**A**) Representative chromogenic RNA-ISH images of *Lta*, (**B**) quantitation of *Lta* area, and (**C**) qPCR for *Lta* in small intestine and colon sections 40h after tamoxifen treatment in mice with the indicated genotype; each data point represents one mouse (mean ± SEM). (**D,F**) Weight curve and (**E,G**) Kaplan-Meier survival curves of the indicated genotypes of tamoxifen-treated mice treated with anti-LTα or isotype control. For panel (**B**), statistical significance was assessed using two-way ANOVA. For panel (**C**), significance was assessed using one-way ANOVA with Dunnett’s multiple comparison test relative to *A20/Abin-1*^*T-ΔIEC*^*Tnf*^−/−^ mice. For panels (**D**) and (**F**), significance was assessed by two-way ANOVA with Bonferroni’s multiple comparison test. For panels (**E**) and (**G**), significance was assessed by Log-rank Mantel-Cox test. Only significant differences are shown; *=p<0.05, **=p<0.01, ***=p < 0.001; ****=p< 0.0001. Scale bars, 50μm. Data represent at least two independent experiments.

To further understand if blocking LTα provided a survival benefit in these mice, we administered a monoclonal blocking antibody against LTα to *A20/Abin-1*^*T-ΔIEC*^*Tnf*^−/−^*MyD88*^+/+^ mice and observed a very small but statistically significant increase in weight and median survival as compared to isotype control, with survival increasing from 4.5 to 5.0 days (**Figure 7D,E**). As we observed a partial improvement in survival in our *A20/Abin-1*^*T-ΔIEC*^*Tnf*^−/−^*MyD88*^+/−^ mice, we administered anti-LTα to these mice and observed a statistically significant improvement in their weight curve, and an increase in median survival from 4.0 to 9.0 days (**Figure 7F,G**). Blocking LTα *in vivo* reduces weight loss and improves survival, particularly when combined with partial deletion of *MyD88*, supporting a model where both microbial signals and LTα contribute to TNF-independent intestinal injury in A20 and ABIN-1-deficient intestinal epithelium.

## DISCUSSION

Genome-wide association studies have implicated hundreds of genes in CD and UC pathogenesis, but it is unclear how these genes increase disease susceptibility or influence treatment response (73). Since IBD is primarily a complex polygenic disease, except for rare monogenic causes, it follows that model systems often incorporate multiple genes simultaneously to decipher these complex epistatic genetic interactions. While we previously showed that A20 and ABIN-1 cooperatively restrict TNF-dependent IEC injury, this study demonstrates that A20 and ABIN-1 preserve intestinal homeostasis and IEC survival by restricting TNF-independent inflammatory injury. Microbial signals, MyD88 activation, and LTα all contribute to TNF-independent intestinal inflammation in the setting of acute A20 and ABIN-1 deletion in IECs. IEC death induced by A20 and ABIN-1 deficiency can also be blocked by simultaneous inhibition of CASP8-dependent apoptosis and RIPK3-dependent necroptosis. Given that A20 and ABIN-1 restrict both TNF-dependent and TNF-independent IEC injury, this study further reveals why these proteins have such potent anti-inflammatory functions in the intestine. Although many previous studies focus on the role of A20 and ABIN-1 in hematopoietic cells, this study and others add to our understanding of the important role these proteins play in non-hematopoietic cells to preserve tissue integrity (9, 16, 38, 40).

Combined acute deletion of A20 and ABIN-1 in the intestinal epithelium is unique in both severity and TNF-independence when compared to deletion of other genes in IECs. For example, the fatal enteritis and colitis that develops after simultaneous deletion of A20 and ABIN-1 in IECs exceeds the severity reported for intestinal epithelial deletion of NEMO (45, 74), FADD (46, 75), CASP8 (46, 72), RIPK1 (43, 76), ATG16L1 (8), or combined deletion of XBP1 and ATG16L1 (7). Acute deletion of SETDB1 (77) in IECs induces mouse lethality with similar kinetics to the combined deletion of A20 and ABIN-1, although SETDB1^−/−^ IECs die primarily from necroptotic cell death. The TNF-independent IEC death observed in *A20/Abin-1*^*T-ΔIEC*^*Tnf*^−/−^ mice is also unusual when compared to other models. The colitis in mice with conditional IEC knock-out of NEMO, FADD, CASP8, and RIPK1 mice is largely reversed by TNF or TNFR1 deletion (43, 45, 46, 74–76). Similarly, the ileitis induced after knock-out of XBP1 in IECs is rescued by TNFR1 deficiency (7). Among these models, deletion of A20 and ABIN-1 is unique in that TNF deletion does not confer any significant survival benefit, and it does not appreciably reduce intestinal injury.

The ileal enteropathy associated with IEC knock-out of NEMO, FADD, or CASP8 is TNFR1-independent, even though colitis in those mice is TNFR1-dependent. Those models provide some perspective on the TNF-independent IEC death observed in *A20/Abin-1*^*T-ΔIEC*^*Tnf*^−/−^ mice. Paneth cell loss after IEC knock-out of FADD and CASP8 is partially reduced by TNFR1 deletion but is further reduced by deletion of Z-DNA binding protein 1 (ZBP1), suggesting that ZBP1 contributes to TNFR1-independent death in the ileum (46). ZBP1 is an interferon-inducible gene product that interacts with RIPK3 via its RHIM domain and activates necroptotic cell death in response to type I & II interferons (78) and viral nucleic acids (79). It is possible that ZBP1 contributes to TNF-independent IEC death after deletion of A20 and ABIN-1, especially since deletion of TRIF only modestly increased median survival. However, ZBP1 has been shown to activate RIPK3 in IECs when FADD-CASP8-apoptotic death is inhibited (46), or in response to RNA from reactivated endogenous retroviruses after SETDB1 deletion (77). In *A20/Abin-1*^*T-ΔIEC*^*Tnf*^−/−^ IECs, both FADD and CASP8 are present to activate apoptotic death, and CC3 is detected in the intestinal epithelium and in intestinal organoids, favoring a central role for CASP8-dependent apoptosis rather than potential ZBP1-dependent necroptosis as the dominant TNF-independent death pathway. In *A20/Abin-1*^*T-ΔIEC*^*Tnf*^−/−^ IECs, RIPK1 is also present and would be predicted to inhibit ZBP1-mediated activation of RIPK3 (80). We cannot entirely exclude a role for ZBP1 in TNF-independent IEC death after deletion of A20 and ABIN-1, but the dominant death pathway in *A20/Abin-1*^*T-ΔIEC*^*Tnf*^−/−^ mice is most consistent with CASP8-mediated apoptosis.

Gasdermin D (GSDMD)-mediated pyroptotic death is another potential TNF-independent death pathway to consider in A20 and ABIN-1 deficient IECs. A20 restricts NLRP3 inflammasome activation (35, 81), so increased susceptibility to pyroptotic death is a possible sequela of acute simultaneous A20 and ABIN-1 deletion in IECs. Canonical inflammasome activation culminates in cleaved caspase-1 and non-canonical inflammasome activation culminates in cleaved caspase-11 in mice (82). Both cleaved caspase-1 and −11 can cleave GSDMD, which in turn causes pyroptotic death (83, 84). Caspase-1 or −11 cleavage of GSDMD could theoretically precipitate pyroptotic death in *A20/Abin-1*^*T-ΔIEC*^*Tnf*^−/−^ mice. Combined deletion of RIPK3 and CASP8 completely rescues A20 and ABIN-1-deficient IECs from death, even when caspase-1 and-11 are present, but RIPK3 regulates inflammasome activation independent of its role in necroptotic death (76). Similarly, the ileitis induced after knock-out of XBP1 in IECs is rescued by TNFR1 deficiency (7). Among these models, deletion of A20 and ABIN-1 is unique in that TNF deletion does not confer any significant survival benefit, and it does not appreciably reduce intestinal injury.

The ileal enteropathy associated with IEC knock-out of NEMO, FADD, or CASP8 is TNFR1-independent, even though colitis in those mice is TNFR1-dependent. Those models provide some perspective on the TNF-independent IEC death observed in *A20/Abin-1*^*T-ΔIEC*^*Tnf*^−/−^ mice. Paneth cell loss after IEC knock-out of FADD and CASP8 is partially reduced by TNFR1 deletion but is further reduced by deletion of Z-DNA binding protein 1 (ZBP1), suggesting that ZBP1 contributes to TNFR1-independent death in the ileum (46). ZBP1 is an interferon-inducible gene product that interacts with RIPK3 via its RHIM domain and activates necroptotic cell death in response to type I & II interferons (78) and viral nucleic acids (79). It is possible that ZBP1 contributes to TNF-independent IEC death after deletion of A20 and ABIN-1, especially since deletion of TRIF only modestly increased median survival. However, ZBP1 has been shown to activate RIPK3 in IECs when FADD-CASP8-apoptotic death is inhibited (46), or in response to RNA from reactivated endogenous retroviruses after SETDB1 deletion (77). In *A20/Abin-1*^*T-ΔIEC*^*Tnf*^−/−^ IECs, both FADD and CASP8 are present to activate apoptotic death, and CC3 is detected in the intestinal epithelium and in intestinal organoids, favoring a central role for CASP8-dependent apoptosis rather than potential ZBP1-dependent necroptosis as the dominant TNF-independent death pathway. In *A20/Abin-1*^*T-ΔIEC*^*Tnf*^−/−^ IECs, RIPK1 is also present and would be predicted to inhibit ZBP1-mediated activation of RIPK3 (80). We cannot entirely exclude a role for ZBP1 in TNF-independent IEC death after deletion of A20 and ABIN-1, but the dominant death pathway in *A20/Abin-1*^*T-ΔIEC*^*Tnf*^−/−^ mice is most consistent with CASP8-mediated apoptosis.

Gasdermin D (GSDMD)-mediated pyroptotic death is another potential TNF-independent death pathway to consider in A20 and ABIN-1 deficient IECs. A20 restricts NLRP3 inflammasome activation (35, 81), so increased susceptibility to pyroptotic death is a possible sequela of acute simultaneous A20 and ABIN-1 deletion in IECs. Canonical inflammasome activation culminates in cleaved caspase-1 and non-canonical inflammasome activation culminates in cleaved caspase-11 in mice (82). Both cleaved caspase-1 and −11 can cleave GSDMD, which in turn causes pyroptotic death (83, 84). Caspase-1 or −11 cleavage of GSDMD could theoretically precipitate pyroptotic death in *A20/Abin-1*^*T-ΔIEC*^*Tnf*^−/−^ mice. Combined deletion of RIPK3 and CASP8 completely rescues A20 and ABIN-1-deficient IECs from death, even when caspase-1 and-11 are present, but RIPK3 regulates inflammasome activation independent of its role in necroptotic death (76). Similarly, the ileitis induced after knock-out of XBP1 in IECs is rescued by TNFR1 deficiency (7). Among these models, deletion of A20 and ABIN-1 is unique in that TNF deletion does not confer any significant survival benefit, and it does not appreciably reduce intestinal injury.

The ileal enteropathy associated with IEC knock-out of NEMO, FADD, or CASP8 is TNFR1-independent, even though colitis in those mice is TNFR1-dependent. Those models provide some perspective on the TNF-independent IEC death observed in *A20/Abin-1*^*T-ΔIEC*^*Tnf*^−/−^ mice. Paneth cell loss after IEC knock-out of FADD and CASP8 is partially reduced by TNFR1 deletion but is further reduced by deletion of Z-DNA binding protein 1 (ZBP1), suggesting that ZBP1 contributes to TNFR1-independent death in the ileum (46). ZBP1 is an interferon-inducible gene product that interacts with RIPK3 via its RHIM domain and activates necroptotic cell death in response to type I & II interferons (78) and viral nucleic acids (79). It is possible that ZBP1 contributes to TNF-independent IEC death after deletion of A20 and ABIN-1, especially since deletion of TRIF only modestly increased median survival. However, ZBP1 has been shown to activate RIPK3 in IECs when FADD-CASP8-apoptotic death is inhibited (46), or in response to RNA from reactivated endogenous retroviruses after SETDB1 deletion (77). In *A20/Abin-1*^*T-ΔIEC*^*Tnf*^−/−^ IECs, both FADD and CASP8 are present to activate apoptotic death, and CC3 is detected in the intestinal epithelium and in intestinal organoids, favoring a central role for CASP8-dependent apoptosis rather than potential ZBP1-dependent necroptosis as the dominant TNF-independent death pathway. In *A20/Abin-1*^*T-ΔIEC*^*Tnf*^−/−^ IECs, RIPK1 is also present and would be predicted to inhibit ZBP1-mediated activation of RIPK3 (80). We cannot entirely exclude a role for ZBP1 in TNF-independent IEC death after deletion of A20 and ABIN-1, but the dominant death pathway in *A20/Abin-1*^*T-ΔIEC*^*Tnf*^−/−^ mice is most consistent with CASP8-mediated apoptosis.

Gasdermin D (GSDMD)-mediated pyroptotic death is another potential TNF-independent death pathway to consider in A20 and ABIN-1 deficient IECs. A20 restricts NLRP3 inflammasome activation (35, 81), so increased susceptibility to pyroptotic death is a possible sequela of acute simultaneous A20 and ABIN-1 deletion in IECs. Canonical inflammasome activation culminates in cleaved caspase-1 and non-canonical inflammasome activation culminates in cleaved caspase-11 in mice (82). Both cleaved caspase-1 and −11 can cleave GSDMD, which in turn causes pyroptotic death (83, 84). Caspase-1 or −11 cleavage of GSDMD could theoretically precipitate pyroptotic death in *A20/Abin-1*^*T-ΔIEC*^*Tnf*^−/−^ mice. Combined deletion of RIPK3 and CASP8 completely rescues A20 and ABIN-1-deficient IECs from death, even when caspase-1 and-11 are present, but RIPK3 regulates inflammasome activation independent of its role in necroptotic death (35, 85, 86), and CASP8 has been reported to mediate GSDMD cleavage under certain conditions (87, 88). This raises the possibility that combined deletion of RIPK3 and CASP8 rescues *A20/Abin-1*^*T-ΔIEC*^ mice due to simultaneous blockade of apoptotic, necroptotic, and pyroptotic death. However, GSDMD-dependent death has been reported in IECs primarily when either FADD-deficiency or catalytically inactive Casp8 are combined with MLKL deficiency (46, 89, 90). Those recent studies suggest that IECs are driven more toward pyroptotic death when apoptosis and necroptosis are inhibited. A central role for pyroptosis therefore would be less likely in *A20/Abin-1*^*T-ΔIEC*^*Tnf*^−/−^ mice, where FADD, active CASP8, RIPK1, RIPK3, and MLKL are all present. Taken together, it is possible that some amount of GSDMD-mediated pyroptotic damage occurs in parallel to apoptotic and necroptotic IEC death in *A20/Abin-1*^*T-ΔIEC*^*Tnf*^−/−^ mice, but apoptotic death is the primary death pathway.

There are multiple potential translational implications for the findings that LTα_3_ and MYD88 contribute to TNF-independent intestinal damage in this model of severe enterocolitis. Deletion of A20 and ABIN-1 unveils a role for LTα_3_-induced TNF-independent IEC death in primary cells both *in vivo* and *in vitro*. To our knowledge, this is the first description of TNF-independent LTα_3_-induced cytotoxicity in IECs in the absence of death-sensitizing agents. There are many potential mechanisms by which patients with IBD may fail to respond to anti-TNF therapy, including activation of alternative inflammatory pathways, neutralizing antibodies, subtherapeutic levels, or inability to block autocrine IEC-derived TNF, but it is tempting to consider whether LTα_3_-mediated TNFR1-induced IEC death could play a role in a subset of patients. Polymorphisms in *LTA* were associated with anti-TNF non-response in a small cohort of patients (14), but this was not reproduced in a larger cohort (91). Moreover, Etanercept, a soluble TNFR2-Fc receptor which blocks TNF and LTα_3_ was ineffective for CD (92). On the other hand, subsequent data suggests that Etanercept is ineffective in part because it does not induce antibody-dependent cell-mediated cytotoxicity or apoptosis of pathogenic TNF-expressing inflammatory cells to the same extent as anti-TNF antibodies (93, 94). Intriguingly, the combination of an anti-TNF monoclonal antibody to neutralize TNF plus Etanercept which additionally neutralizes LTα_3_ was effective in a case report of a patient with severe HLA-B27-associated arthropathy who had failed treatment with either agent alone (95). The other TNF-independent injury pathway highlighted in this study is MYD88. Microbial signals activating MYD88 in the hematopoietic compartment contribute to intestinal injury, but it would be instructive to understand which microbes are most pathogenic in *A20/Abin-1*^*T-ΔIEC*^*Tnf*^−/−^ gnotobiotic mice. Interestingly, the fact that some germ-free *A20/Abin-1*^*T-ΔIEC*^*Tnf*^−/−^ mice die suggests that sterile inflammation is sufficient to induce death. MYD88 mediates signaling downstream of IL-1β and IL-18 which could potentially explain the death of some germ-free *A20/Abin-1*^*T-ΔIEC*^*Tnf*^−/−^ mice. Both IL-1β and IL-18 have been proposed as potential mediators of anti-TNF non-response in patients with IBD (96–98). Ultimately, having animal models to investigate TNF-independent intestinal injury could prove useful in understanding anti-TNF non-response in subsets of patients.

In summary, we discovered that microbial signals, MYD88 signaling, and LTα_3_ drive severe TNF-independent enterocolitis after acute deletion of two IBD-associated genes *A20* and *Abin-1*, which can be prevented by combined deletion of *Ripk3* and *Casp8.* This study highlights that A20 and ABIN-1 cooperatively maintain intestinal homeostasis by inhibiting both TNF-dependent and TNF-independent inflammatory intestinal injury. Understanding the genetic determinants of intestinal epithelial health, their epistatic relationships, and how they could influence therapeutic response in models of IBD will provide important mechanistic insights to inform future translational studies.

## Methods

### Mice

*A20*^*fl*^ and *Abin-1*^*fl*^ mice were generated in the Ma laboratory and were described previously (9, 28, 62, 99). *Tnf*^−/−^ mice were purchased from Jackson Laboratories (Bar Harbor, MA), X. Wang provided *Ripk3*^−/−^ mice, and transgenic mice with a tamoxifen-inducible Cre recombinase under the control of the villin-promoter (*Vil-cre-ER*^*T2*+^) mice were a kind gift from S. Robine (Institut Curie-CNRS, Paris, France) (100). These alleles were backcrossed to *A20*^*fl*^ and *Abin-1*^*fl*^ mice transgenic mice for > 8 generations as previously described (9). Acute deletion of floxed *A20* and *Abin-1* exons *in vivo* by oral gavage was performed using tamoxifen (2 mg/day, Sigma, T5648) for 3 consecutive days as indicated for survival analysis. *MyD88*^+/−^ and *Trif*^+/−^ breeders were maintained on antibiotic water *ad libitum* with ampicillin 1mg/ml (Sigma), vancomycin 0.5mg/ml (Sigma), and neomycin 1mg/ml (Sigma), and offspring were transitioned to regular water at weaning. For *in vivo* monoclonal antibody treatment, mice were injected with 370μg of antibody on days −2, 0, 1, 2, 5, 7, and 9. For all experimental mice, genotypes were confirmed twice. When possible, littermates were used as controls. Both males and females were included in all experiments with no observable gender differences. Mice were analyzed between 7-12 weeks of age for all experiments.

### Germ-free derivation, maintenance, and conventionalization

Germ-free A20/Abin-1^T-*ΔIEC*^*Tnf*^−/−^ mice were derived by Cesarean delivery into the UCSF Gnotobiotic Core Facility (gnotobiotics.ucsf.edu) and maintained in germ-free isolators. Stool pellets from isolators in the Gnotobiotic facility were screened every 2-3 weeks by culture and qPCR, whenever mice were transferred from breeding to experimental isolators, and at the beginning and end of each experiment. 515F and 806R universal primers were used to amplify the V4 region: 515F-GTGCCAGCMGCCGCGGTAA; 806R-GGACTACHVGGGTWTCTAAT. For cecal content conventionalization experiments, cecal contents from specific-pathogen free (SPF) mice were resuspended in 10% weight/volume (w/v) Bacto Brain Heart Infusion (BHI) (VWR) supplemented with 0.05% w/v L-cysteine (Sigma) in aerobic conditions, filtered over a 100μm filter, and 200μl was administered to germ-free mice by oral gavage 5-7 days prior to tamoxifen treatment.

### CRISPR/Cas9 editing of *MyD88*, *Trif*, and *Casp8*

Two guide RNAs for each target gene were designed using Benchling (Benchling, San Francisco, CA). The following guide RNAs (gRNAs) were selected for editing A20/Abin-1^T-*ΔIEC*^*Tnf*^−/−^ mice: *MyD88* guide 1 CCCACGTTAAGCGCGACCAA and *MyD88* guide 2 AAGGAGCTGAAGTCGCGCAT targeting exon 1 of *MyD88*; *TRIF* guide 1 TCTGGAACGCTAATTTCGTG and *TRIF* guide 2 CAAGCTATGTAACACACCGC targeting exon 2 of *Trif*. The following gRNAs were selected for editing A20/Abin-1^T-*ΔIEC*^*Ripk3*^−/−^ mice: *Casp8* guide 1 TAGCTTCTGGGCATCCTCGA and *Casp8* guide 2 CTTCCTAGACTGCAACCGAG targeting exon 2 of *Casp8*. The corresponding crRNAs targeting each gene and tracrRNA were annealed and ribonucleoprotein (RNP) complexes were prepared with HiFi Cas9 nuclease ribonucleoprotein (IDT) according to manufacturer instructions. RNPs were electroporated into mouse zygotes as previously described (52), with the main exception that the standard square curve electroporation (two pulses at 30V for 3 ms with an interval of 100 ms) was performed once or four times with an interval of 3s Gene Pulser Xcell (Biorad). Zygotes were cultured to two-cell embryos and transferred into pseudopregnant females. All mutations were confirmed by cloning and sequencing. Mice with a confirmed deletion or premature stop codon were then backcrossed 3 generations to the parental strains to breed out off-target edits before analysis.

### Intestinal Epithelial Cell Enteroid Culture and Confocal Microscopy

For enteroid culture, intestinal crypts were isolated from the small intestine as previously described(101), with the modifications of substituting 10% R-spondin-1 conditioned medium for recombinant R-spondin1, and the addition of Normocin (100 mg/ml, Invivogen). R-spondin1 expressing 293T cells were a kind gift from Dr. Noah Shroyer (Baylor College of Medicine). For all enteroid experiments, enteroids were derived from at least 2 mice on separate occasions and representative data are shown. Deletion of *A20* or *Abin-1 in vitro* was performed via treatment with 4-OHT (200nM, Sigma). Confocal imaging of enteroids was performed on a Leica SP5 laser scanning confocal system (Leica Microsystems) using a 10X dry objective. Images were acquired in a format of 512×512, with a line average of at least 3, scan speed of 400 Hz, and pinhole airy unit 1. Excitation for both PI and brightfield was done with the 488nm laser line at 30% power with a detection band of 550-732nm. Image analysis was performed on the Leica Application Suite (Leica Microsystems).

### Cell Death Assays

Enteroid death assays were performed by resuspending in Matrigel (Corning) and plating 25μl per well in 96-well flat-bottom opaque plates (Nunc). After 24h, enteroids were stimulated as indicated to a final volume of 200μl. Viability was measured using the CellTiter Glo 3D assay (Promega) according to the manufacturer’s specifications, with the exception that 100μl of reagent was added to 200μl of culture for a final volume of 300μl prior to reading. Luminescence was read on a SpectraMax M5® (Molecular Devices), analyzed using SoftMax Pro (Molecular Devices), and expressed as percent viability relative to vehicle control.

### Cell Signaling Assays and Immunoblot Analysis

For enteroid lysates, cultures were resuspended in Cell Recovery Solution (Corning) supplemented with 10μM Y-27632 (Calbiochem) and incubated for 15min on ice, followed by centrifugation at 500xg for 5min. Cell pellets were lysed in ice-cold NP40 lysis buffer (1% NP40 v/v, 50mM Tris HCl pH 7.4, 150mM NaCl, and 10% glycerol v/v) supplemented with complete EDTA-Free Protease Inhibitor Cocktail (Roche), phosphatase inhibitors (1mM Na3VO4 and 10mM NaF), and 10mM N-ethylmaleimide. After lysis, samples were centrifuged for 20min at 21,130xg to remove debris, and the supernatants were quantitated using the BCA Protein Assay Kit (Pierce). Lysates were normalized and denatured in LDS Sample Buffer (Invitrogen), followed by resolution on NuPage precast 4-12% Bis-Tris gels (Invitrogen) and transferred to PVDF for immunoblotting.

### qPCR

1cm of small intestine or colon was flushed with saline solution and placed in RNAlater (ThermoFisher Scientific) and stored at 4C. The following day, the solution was aspirated, and samples were stored at −70C. Samples were then thawed and homogenized in Buffer RLT (Qiagen) supplemented with β-mercaptoethanol using Lysing Matrix D tubes (MP Biomedicals) on the FastPrep-24 (MP Biomedicals), followed by QIAshredder homogenization (Qiagen). RNA was prepared using the RNeasy Mini Kit (Qiagen) with on-column DNase digestion according to manufacturer’s instructions. Quality was confirmed using the NanoDrop ND-1000 (ThermoFisher Scientific) and Agilent RNA 6000 Nano Kit on the Agilent 2100 Bioanalyzer (Agilent), according to manufacturer’s instructions. cDNA was synthesized from 2000ng of total RNA using the High-Capacity RNA-to-cDNA kit (Thermo Scientific). qPCR was performed using TaqMan probes for mouse *Lta* (Mm00440228_gH) and mouse *Actb* (Mm02619580_g1), and the TaqMan® Universal Master Mix II with UNG on the QuantStudio 6 (ThermoFisher Scientific). Relative gene abundance was normalized to the mean expression of the house-keeping gene *Actb*, and 2^−ΔΔCt^ was calculated relative to *A20*^*fl/fl*^*Abin-1*^*fl/fl*^*Tnf*^−/−^. All samples were run in duplicate. Data were analyzed using QuantStudio RealTime PCR Software (Thermo Scientific).

### Histology and Immunohistochemistry

Mice were individually euthanized, dissected, and their resected intestinal tissue was immediately fixed in 10% neutral-buffered formalin (Sigma) 40h after treatment with tamoxifen. Tissue specimens were fixed for 16-24 hours, washed in PBS three times, then placed in 70% ethanol and shipped to Histowiz (Brooklyn, NY) for paraffin embedding, sectioning, staining, and high-resolution (40X) scanning according to standard protocols. H&E histologic severity quantitation was performed using a scoring system previously described for genetic IBD models affecting the small intestine (score range 0-15) or colon (score range 0-12) (102). For CC3-stained sections (Cell Signaling Technology 9661), 10 consecutive crypts from each of 5 sections for small intestine and colon (50 crypts per mouse per segment) were counted. Chromogenic RNA *in situ hybridization* (RNA-ISH) was performed using RNAscope^®^ Probe- Mm-*Lta* (Advanced Cell Diagnostics 317231) and RNAscope 2.5 HD Assay- Brown Kit (Advanced Cell Diagnostics 322370) according to the manufacturer’s specifications. Slides were immediately covered with a coverslip using Cytoseal. Slides were then scanned at 40X magnification with a Zeiss Axio ScanZ.1 (Zeiss). DAB-positive areas were identified against a hematoxylin background in all 5 sections of small intestine and colon on each slide using QuPath v0.2.3. At least two and up to three blinded readers assessed histologic disease severity, CC3 quantitation, and *Lta* quantitation and the results were averaged.

### Antibodies and reagents

Antibodies directed against A20 (Cell Signaling Technology, 5630), ABIN-1 (Sigma HPA037893), Caspase 8 (Cell Signaling Technology mouse specific 4927 and 4790), CC8 (Cell Signaling Technology 8592), RIPK1 (Cell Signaling Technology 3493), Caspase 3 (Cell Signaling Technology 9662), CC3 (Cell Signaling Technology 9661), PARP (Cell Signaling Technology 9542), mouse RIPK3 (Cell Signaling Technology 95702), phospho-RIPK3 (T231/S232) (Cell Signaling Technology 91702), phospho-RIPK1 (Ser166) rodent specific (Cell Signaling Technology 31122), and GAPDH (Millipore MAB374), were used for western blot. Anti-CC3 (Cell Signaling Technology 9661) was also used for IHC studies. Mouse IFN-β ELISA (PBL assay bioscience) was used according to the manufacturer’s instructions. Recombinant mouse TNF, mouse IL-18, mouse TL1A, mouse LTα_1_β_2_, human LTα_3_, mouse TNFR1-Fc, and mouse LTBR-Fc were purchased from R&D systems.

Recombinant mouse RANKL was purchased from BioLegend. Recombinant mouse IL-1β and mouse LIGHT were purchased from Peprotech. Pam3CSK4, poly(I:C) HMW, and LPS from E. coli O111:B4 were purchased from Invivogen. Necrostatin-1s (BioVision 2263) and Emricasan (MedChem Express HY-10396) were used at 50μM final concentration. S5H3.2.2 hybridoma cell line (ATCC) expressing monoclonal hamster anti-mouse LTα (66) was provided to BioXcell (Lebanon, New Hampshire) for production of purified monoclonal antibody for *in vivo* use. *InVivo*MAb Armenian hamster IgG isotype control anti-glutathione S-transferase (BioXcell #BE0260) was used as a negative control.

### LTα_3_-Fc conditioned media

Codon-optimized gblocks (IDT) encoding mouse LTα (amino acids 1-202; UniProtKB/Swiss-Prot P09225.1) followed by a GSG linker and mouse IgG2a Fc fusion (amino acid 99-330; UniProtKB/Swiss-Prot P01863) were synthesized and then cloned into pENTR (Addgene Plasmid #17398) using InFusion cloning according to manufacturer’s instructions (Clontech). The insert sequence was confirmed and then cloned into pLEX_307 (Addgene Plasmid #41392) using the Gateway Cloning LR reaction according to manufacturer’s instructions (ThermoFisher Scientific). Lentivirus particles were prepared by co-transfection with the packaging plasmid psPAX2 (Addgene Plasmid #12260) and the envelope plasmid pMD2.G (Addgene Plasmid #12259) into HEK293T (ATCC) cells using Lipofectamine 2000 (ThermoFisher Scientific). The lentivirus-containing medium was collected at 72 h post-transfection. After centrifugation, the lentivirus medium was filtered using a 0.45 μm syringe filter. HEK293T cells were transduced with diluted lentivirus and selected with puromycin 5 ug/ml for several passages. The puromycin was removed and mLTα-Fc conditioned media (mLTα-Fc CM) was collected and filtered over 0.22μm filter. The mLTα-Fc CM was quantitated by ELISA with goat polyclonal LTα capture antibody (R&D systems AF749), rat monoclonal LTα detection antibody (R&D systems MAB749) biotinylated with ChromaLink biotin according to manufacturer’s instructions (Trilink Biotechnologies), a recombinant mouse lymphotoxin alpha ELISA reference (a gift from R&D systems), peroxidase streptavidin (Jackson ImmunoResearch), and 1-Step Ultra TMB-ELISA (ThermoFisher Scientific).

### Statistics

Statistical analysis was performed with Graphpad Prism 9 (Graphpad Software, San Diego, CA). Kaplan-Meier survival curve comparisons were performed using the log-rank Mantel-Cox test. Comparisons between two groups were performed by two-tailed unpaired Student’s t-test. Multigroup comparisons were performed by one-way analysis of variance (ANOVA) if we were comparing one variable per group or two-way ANOVA if there were multiple variables per group. When comparing every mean to every other mean by ANOVA, Tukey’s multiple comparison test was used. When comparing each mean to a control, Dunnett’s multiple comparison test was used. When comparing only a subset of means, Bonferroni’s multiple comparison test was used. p< 0.05 was used as the threshold for statistical significance. All experiments shown represent at least two independent repetitions.

### Study approval

All animal studies were conducted in accordance with the University of California, San Francisco Institutional Animal Care and Use Committee (AN183350).

## Author Contributions

M.G.K. and A. Ma conceived of the project. M.G.K. oversaw the project and designed and performed the experiments with assistance from I.R and E.M. I.R. performed *in vivo* experiments, *in vitro* death assays, and helped generate LTα_3_-Fc CM. Co-first author order determined by duration of time spent on the project. E.M. performed enteroid immunoblot analysis and *in vitro* assays. E.M. and I.R. performed RNA-ISH analysis with advice and guidance from O.D.K. Z.L. performed CRISPR/Cas9 zygote editing with guidance from A. Marson. J.L.B., X.S., Y.Y.R, Z.W., R.A., and P.A. assisted with mouse breeding, genotyping, and histology quantitation. K.L., P.J.T., and J.A.T. derived germ-free mice and performed germ-free experiments in the UCSF Gnotobiotics Core Facility. B.R. provided a key observation that lymphotoxin could contribute to TNF-independent death in this model. B.R and L.S. critically reviewed the manuscript and provided helpful discussion. M.G.K performed the statistical analyses and wrote the manuscript with editing from I.R., E.M, B.A.M, A.M., and input from all co-authors.

## Acknowledgements

MGK was supported by a Career Development Award from the Crohn’s and Colitis Foundation and NIH K08 DK123202. Michael George Kattah, M.D., Ph.D., holds a Career Award for Medical Scientists from the Burroughs Wellcome Fund. This work was also supported by R01 NIH grant DK095693 and the Crohn’s and Colitis Foundation (A.Ma). A.Marson holds a Career Award for Medical Scientists from the Burroughs Wellcome Fund, is an investigator at the Chan Zuckerberg Biohub, and is a recipient of The Cancer Research Institute (CRI) Lloyd J. Old STAR grant. The Marson lab has received funds from the Innovative Genomics Institute (IGI), the Simons Foundation, and the Parker Institute for Cancer Immunotherapy (PICI).

## Supplemental Material

**Supplemental Figure 1.**
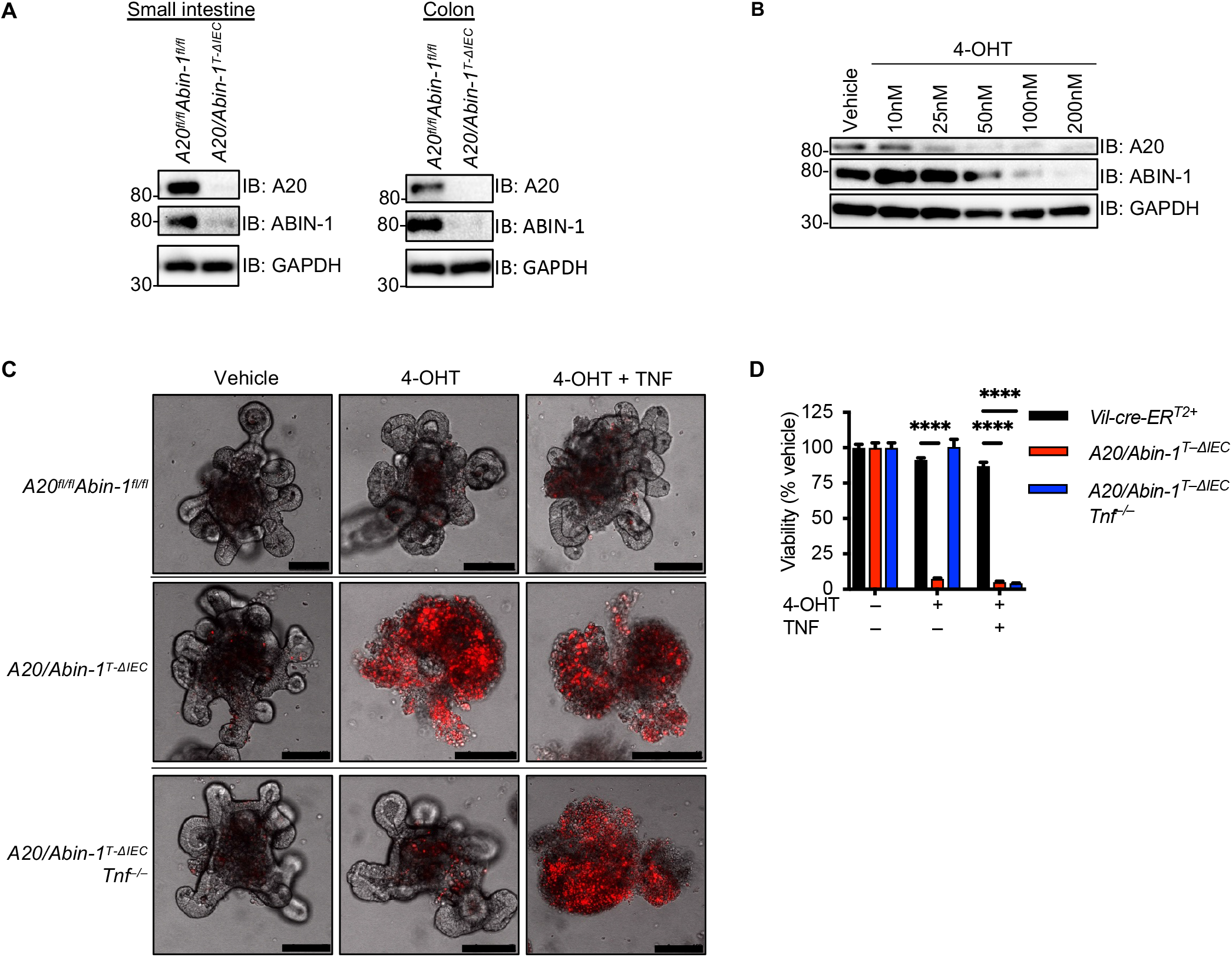
Tamoxifen-induced deletion of floxed A20 and ABIN-1 on a *Vil-cre-ERT*^2+^ background *in vivo* and *in vitro*. (**A**) Immunoblotting analysis of IECs with the indicated genotypes 40h after treatment with 2mg tamoxifen *per os* (*p.o.*). (**B**) Immunoblotting analysis of *A20/Abin-1*^*T-ΔIEC*^*Tnf*^−/−^ enteroid cultures treated with the indicated concentration of 4-OHT for 24h. For panels (**A,B**), lysates were immunoblotted with the antibodies indicated on the right. (**C**) Representative confocal microscopy images of PI-stained enteroids from indicated genotypes of mice treated with vehicle or 4-OHT for 24h followed by 24 h of 2.5 ng/ml TNF as indicated. Scale bars, 100μm. (**D**) Quantitative luminescent cell viability assay of enteroid cultures treated as described in (**C**) (mean ± SEM). For panel (**D**), significance was assessed by two-way ANOVA with Dunnett’s multiple comparison test comparing *A20/Abin-1*^*T-ΔIEC*^ and *A20/Abin-1*^*T-ΔIEC*^*Tnf*^−/−^ to control *Vil-cre-ERT*^2+^ enteroids.

**Supplemental Figure 2.**
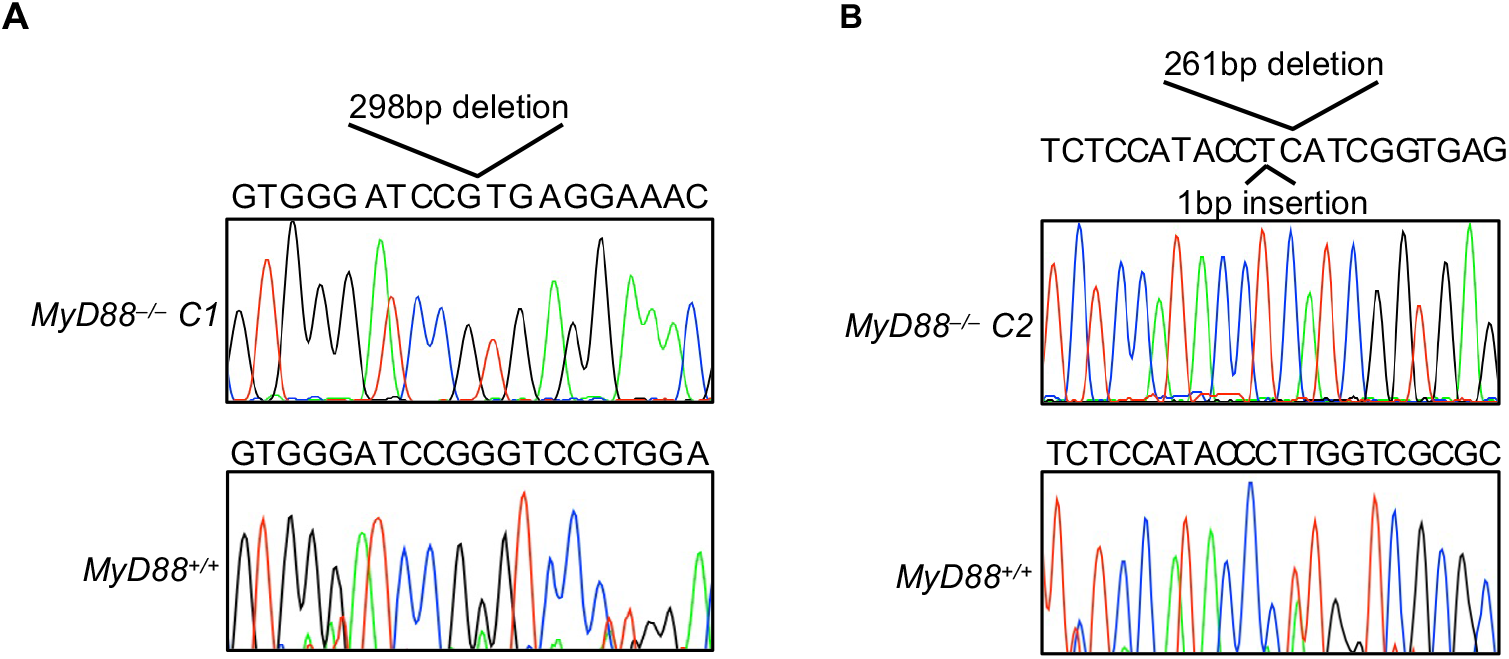
Editing of *MyD88* in *A20/Abin-1*^*T-ΔIEC*^*Tnf*^−/−^ mice. Nucleotide chromatogram and sequence of the *MyD88* PCR amplicon surrounding the CRISPR-Cas9-targeted site for (**A**) *MyD88 C1* and (**B**) *MyD88 C2* in *A20/Abin-1*^*T-ΔIEC*^*Tnf*^−/−^ mice.

**Supplemental Figure 3.**
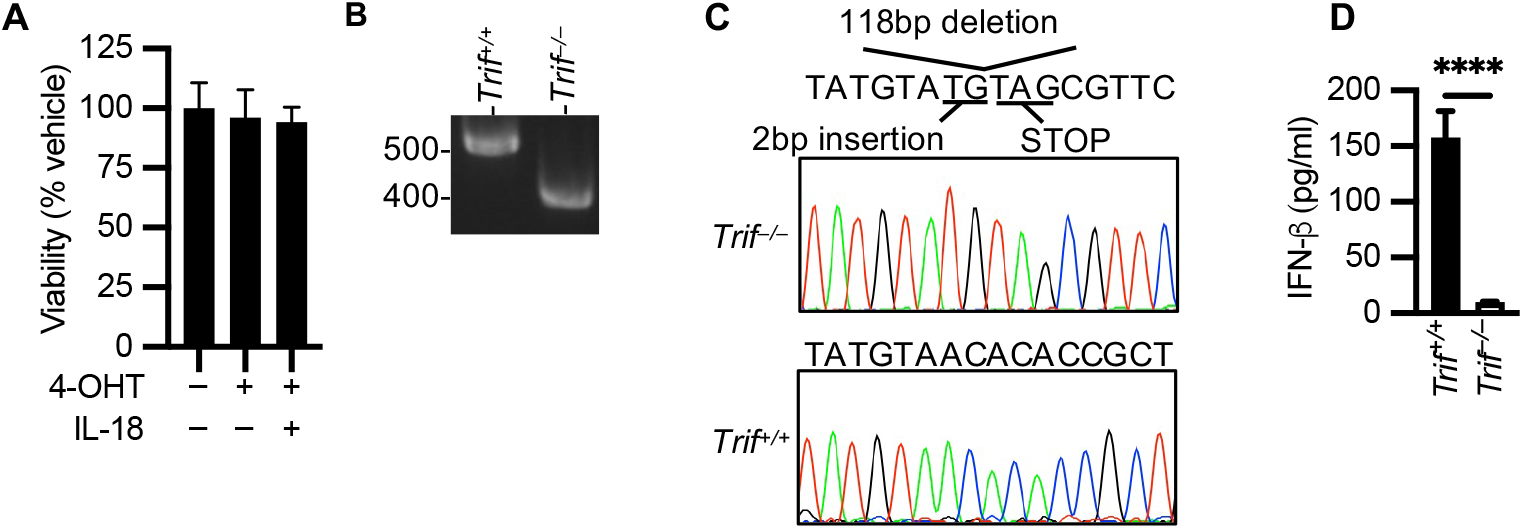
IL18 stimulation of *A20/Abin-1*^*T-ΔIEC*^*Tnf*^−/−^ enteroids and editing of *Trif* in *A20/Abin-1*^*T-ΔIEC*^*Tnf*^−/−^ mice. (**A**) Quantitative luminescent cell viability assay of *A20/Abin-1*^*T-ΔIEC*^*Tnf*^−/−^ enteroid cultures treated with the indicated stimuli (mean ± SEM; IL-1β 25ng/ml). (**B**) Agarose gel electrophoresis and (**C**) nucleotide chromatogram and sequence of the *Trif* PCR amplicon surrounding the CRISPR-Cas9-targeted site in *Trif* for *A20/Abin-1*^*T-ΔIEC*^*Tnf*^−/−^ mice. (**D**) IFN-β ELISA of mouse splenocytes with the indicated genotype stimulated with 100 μg/ml poly(I:C). For panel (**A**) significance was assessed using one-way ANOVA with Dunnett’s multiple comparison test relative to vehicle alone. For panel (**D**), significance was assessed using unpaired t-test. Only significant differences are shown; ****=p< 0.0001. Data represent at least two independent experiments.

**Supplemental Figure 4.**
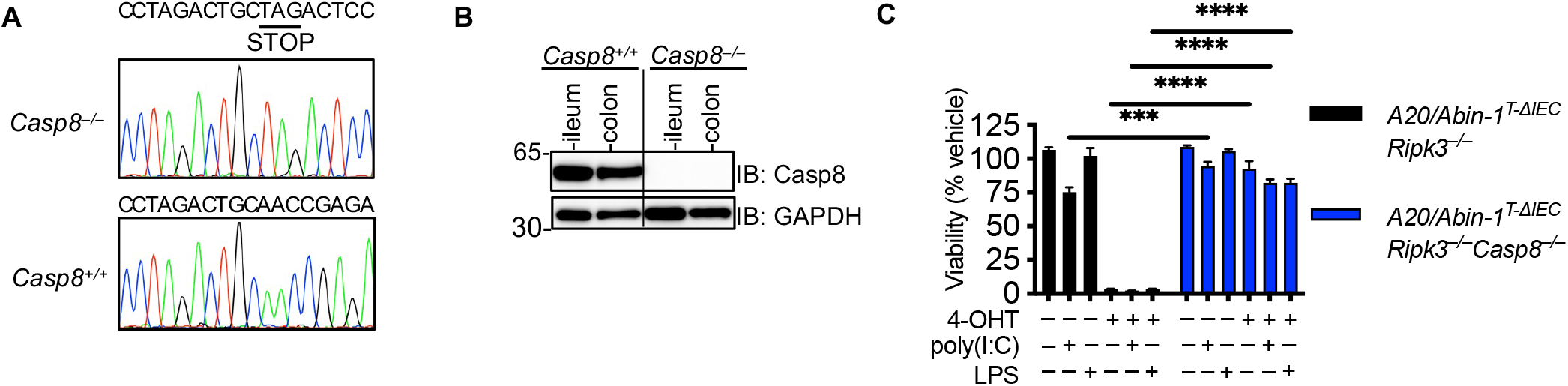
Deletion of *Casp8* in *A20/Abin-1*^*T-ΔIEC*^*Ripk3*^−/−^ mice and poly(I:C) stimulation of *A20/Abin-1*^*T-ΔIEC*^*Ripk3*^−/−^ and *A20/Abin-1*^*T-ΔIEC*^*Ripk3*^−/−^*Casp8*^−/−^ enteroids. (**A**) Nucleotide chromatogram and sequence of the *Casp8* PCR amplicon surrounding the CRISPR-Cas9-targeted site in *Casp8* for *A20/Abin-1*^*T-ΔIEC*^*Ripk3*^−/−^ mice. (**B**) Immunoblot of intestinal epithelial cells from *A20/Abin-1*^*T-ΔIEC*^*RIpk3*^−/−^ mice with the indicated *Casp8* genotypes. (**C**) Quantitative luminescent cell viability assay of enteroids with the indicated genotype treated with indicated stimuli, (mean ± SEM). For panel (**C**) statistical significance was assessed by two-way ANOVA with Bonferroni’s multiple comparison test comparing between genotypes for each stimulation condition. Only significant differences are shown; ***=p < 0.001; ****=p< 0.0001. Data represent at least two independent experiments.

